# GuFi phages represent the most prevalent viral family-level clusters in the human gut microbiome

**DOI:** 10.64898/2026.01.26.701711

**Authors:** Hanrong Chen, Andrew Ting, Indrik Wijaya, Xin An Tang, Yue Yuan On, Jean-Sebastien Gounot, Windsor Koh, Ivan Liachko, Yik-Ying Teo, Henning Seedorf, Kun Qu, Niranjan Nagarajan

## Abstract

Despite being important ecological modulators of the gut microbiome, bacteriophage diversity and function remain under-characterized. We show that short-read metagenomic surveys can miss even globally highly prevalent viral family-level clusters (VFCs), that can be readily assembled and characterized with long-read metagenomic data from a relatively small cohort (n=109). While gut Bacteroidota phages have been the prevailing focus in the literature, we show that highly prevalent gut phage families frequently have Firmicutes hosts (termed GuFi phages), with broad host ranges verified using proximity-ligation (Hi-C) sequencing data. High-throughput sequencing of virus-like particles from fecal samples detected frequent enrichment of GuFi phages across samples, revealing their under-appreciated impact on the gut microbiome. We report the first *in vitro* induction and imaging of members of prevalent GuFi clades including the candidate orders *Heliusvirales*, *Astravirales* (VFC 2) and *Suryavirales* (VFC 4). Our findings underscore the importance of GuFi phages with broad host ranges in the gut microbiome, and the utility of long-read sequencing for viral discovery, paving the way for deeper insights into the role of bacteriophages in human health and disease.

## Introduction

The human gut microbiome is a complex community of bacteria, archaea, viruses, and fungi that play critical roles in health and disease^1,2^. Bacteriophages (phages), viruses that infect bacteria, are present in the human gut at numbers comparable to bacteria but remain a relatively underexplored component of this ecosystem^3,4^. Phages have been shown to influence early-life microbiome assembly^5,6^, colonization by invading strains^7,8^, and horizontal gene transfer following antibiotic perturbation^9^, where their impact extends beyond their immediate hosts to shape broader microbial community dynamics^10^. The gut virome has been associated with human diseases such as inflammatory bowel disease^11^ and asthma^12^, with studies showing effectiveness of fecal virome transplants for *Clostridioides difficile* infections^13^, though the underlying mechanisms remain unclear. A deeper understanding of phage diversity and ecology in the gut is thus important to clarify their function in health and disease, and pave the way for novel phage-based microbiome therapeutics^14,15^.

Metagenomic studies have been critical to uncover the vast diversity of human gut phages, most of which are known solely from sequence data and remain uncultured^16–20^, with vast amounts of newly assembled sequences (>15 million in IMG/VR^20^ alone) leading to a fundamental and ongoing shift from morphology-based to sequence-based viral taxonomy^21^. Major families of gut phages are known to primarily infect members of the core gut bacterial phylum Bacteroidota, including crAssphages^22–24^, Gubaphage^17^, and some of the recently described clades in a Japanese cohort^25^. In comparison, phages infecting other major gut bacterial phyla including Firmicutes, Actinobacteria and Verrucomicrobia are less well-described^25,26^. The ability to recover high-quality and complete viral genomes can substantially impact phage family discovery due to the absence of universal marker genes, the rapid nature of viral evolution, and extensive genetic mosaicism^27,28^. However, nearly all large-scale gut virome surveys to date^16–20,25,29–33^ have relied on short-read sequencing. Long-read metagenomics has been shown to improve the number and completeness of viral genomes recovered^29^, capture structural variation^34^, and elucidate prophage integration dynamics^35,36^, but the potential of long reads for *de novo* discovery of gut viral families has not been explored.

To investigate this, we systematically leveraged a gut microbiome cohort with short and long-read metagenomic sequencing data^37^ (SPMP, n=109) to assemble and annotate >30,000 viral operational taxonomic units (vOTUs) and >300 viral family-level clusters (VFCs). Comparison of short-read and long-read**–**augmented metagenomic assemblies highlighted >2.4× improvement in median vOTU contiguity (>10-fold increase for vOTUs longer than 30kbp), 2× increase in vOTU recovery, and frequent consistent fragmentation of short-read vOTUs affecting viral family-level clustering. Strikingly, >60% of the vOTUs and 11 out of the top 12 VFCs (prevalence >50%) in our cohort have not been described before. Globally, 6 out of the newly identified VFCs are among the most prevalent 10 families, with VFC 3 being the most prevalent (>80%). Intriguingly, 9 out of the top 12 VFCs identified in our study were found to have Firmicutes hosts, with broad host ranges confirmed using extensive Hi-C data (n=108), and more diverse hosts relative to other gut phage families (>4×). Sequencing of virus-like particles (n=64) identified an enrichment of most GuFi phage families across many samples, and prophage induction experiments with diverse Firmicutes strains helped obtain the first transmission electron microscopy images for prevalent GuFi clades including the candidate orders *Heliusvirales*^26^, *Astravirales* (VFC 2) and *Suryavirales* (VFC 4). Our findings underscore the importance of phage families infecting diverse Firmicutes species in the gut microbiome, laying the foundation for understanding their role in gut ecology and health.

## Results

### Long-read metagenomic sequencing substantially improves viral genome recovery

To study the utility of long-read sequencing for viral discovery, we complemented long and short-read gut metagenomic datasets for a cohort (SPMP^37^; n=109), with Hi-C sequencing and virus-like particle sequencing data (**Fig. 1A**; **Methods**). Leveraging the high-quality of the hybrid assemblies generated (accurate short-read contigs, with contiguity boosted using long reads^38^) in terms of contiguity and base-level accuracy^37^, a combination of state-of-the-art viral identification tools^39–41^ was utilized to recover a set of 30,510 non-redundant viral operational taxonomic units (vOTUs at 95% average nucleotide identity over 85% alignment fraction, as species-level proxy for viruses^42^; **Supplementary Fig. 1A-B**; **Methods**; **Supplementary Data 1**). A substantial number of these (n=3,511 vOTUs) were determined to be high-quality (>90% complete) or complete genomes (**Supplementary Fig. 2A**), even though this may be an underestimate as noted from analysis of circular vOTUs (**Supplementary Fig. 2B-C**). In comparison, short-read**–**only assemblies rarely generated vOTUs longer than 30kbp (median length=8.4kbp; >10× reduction relative to long-read**–**augmented assemblies; **Fig. 1B**), with only 258 vOTUs assessed as being high-quality or complete (>13× reduction), and with a 2× reduction in vOTU recovery based on short-read viral contigs^29,43^ (**Supplementary Fig. 3A**). Even among shared vOTUs (n=11,098), long-read**–**based contigs were notably longer (>2.4× median length, **Supplementary Fig. 3B**), even though these contigs were well covered by short reads (median coverage of 99.6%; **Supplementary Fig. 3C**), highlighting the outsized impact of assembly fragmentation on viral genome and vOTU recovery. Comparison of the assembled vOTUs with 11 of the largest and most widely used viral genomic databases^16–20,25,30–34^ highlighted significant novelty (>62% novel; **Fig. 1C**), despite their construction from extensive short-read and limited long-read metagenomic surveys (>6 million vOTUs; **Methods**). The largest individual overlaps were with IMG/VR (32%) and GPD (22%), while the overlap with other gut phage databases typically accounted for <10% of the vOTUs assembled. These results emphasize the substantial utility of long-read sequencing for capturing viral diversity through metagenomic surveys.

**Figure 1.**
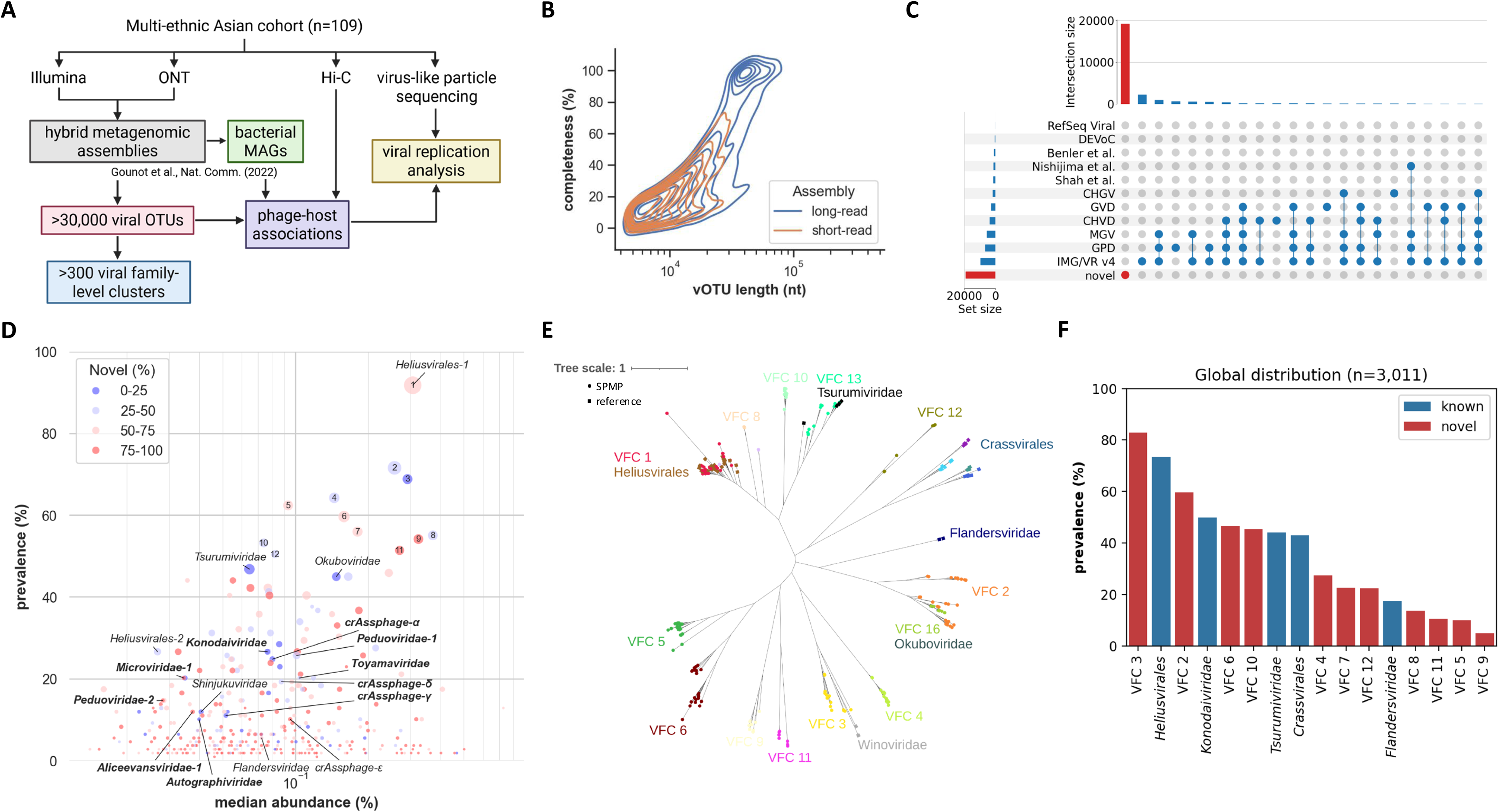
Improved viral genome recovery and discovery of globally prevalent viral family-level clusters via long-read metagenomic sequencing of a multi-ethnic Asian cohort. (A) Schematic of sequencing approaches used to characterize the diversity, host associations, and replication activity of viral genomes in the SPMP cohort (n=109). (B) The addition of long-read sequencing notably increased the number and completeness of viral OTUs (vOTUs) recovered. (C) The vOTUs exhibit substantial novelty with respect to 11 publicly available gut viral genome datasets (>62% novel). (D) Prevalence and median abundance of viral family-level clusters (VFCs), identified through a gene sharing approach applied on the vOTUs and reference genomes (**Methods**). VFCs are color-coded based on the percentage of novel vOTUs, and marker sizes are proportional to VFC size. Reference families are labelled, with ICTV-ratified families in bold. The top 12 VFCs are numbered by prevalence in this cohort. (E) Phylogenetic tree of portal proteins shows distinct clustering of highly prevalent VFCs and several reference families. Proteins sampled from SPMP and reference genomes (**Methods**) are marked in circles and squares, respectively, and color-coded by VFC/reference clade. (F) Global prevalence of VFCs. Short reads from 3,011 publicly available samples were mapped to the vOTUs and reference genomes (**Methods**), finding several VFCs that are highly prevalent not only in the SPMP cohort, but also globally. VFCs corresponding to known viral families are in blue, and other VFCs (>50% prevalence in SPMP cohort) are in red. *Crassvirales* VFCs were grouped together in this analysis.

### Identification of globally prevalent viral family-level clusters using a population-specific long-read metagenomics dataset

While human gut phages are highly individual-specific at the vOTU level^44,45^, prevalent clades at higher taxonomic levels remain largely unmapped despite recent overhauls in viral taxonomy^21,46^. To assess if improvements in long-read metagenomics-based viral OTU recovery could enable discovery of key viral families, the 3,511 high-quality and complete vOTUs were clustered together with publicly available genomes with family-level taxonomic assignments^24,32,25,26,33^, including those officially ratified by the International Committee on Taxonomy of Viruses^47^ (ICTV). As existing viral identification tools do not perform detailed taxonomic classification^39–41^, and other popular tools are not well-suited for *de novo* family-level discovery^48^, a frequently used clustering approach based on sharing of protein families^23,25,49^ was adopted to construct viral family-level clusters (VFCs, n=351 non-singleton; **Supplementary Fig. 4**; **Methods**; **Supplementary Data 2-3**) with broad agreement with known viral families (**Supplementary Fig. 5**). Clustering of short-read vOTUs and reference genomes using the same approach identified far fewer members of even the most prevalent VFCs in the cohort (<20% on average; **Supplementary Fig. 6A**), highlighting the challenges of VFC construction with short-read metagenomic assembly. One potential explanation for this is consistent fragmentation of short-read**–**based vOTUs as was seen for the most prevalent VFC (VFC 1), where the assembled short-read contigs consistently covered <50% of the vOTU (**Supplementary Fig. 6B**), with a consistent break in the middle of the vOTU (**Supplementary Fig. 6C-D**) that was also seen in the cognate sample from which the vOTU sequence was assembled. As the long-read contigs in this study were constructed with short-read contigs as a starting point^38^, they provide a more controlled demonstration of the utility of long-reads for improving assembly contiguity (**Methods**). In addition, the assembled long-read and short-read contigs have similar gene length distributions (though long-read contigs help identify some longer genes; **Supplementary Fig. 6E**), further emphasizing the importance of genomic contiguity derived from long-read assembly for identifying the shared protein families that are critical for VFC construction.

Analysis of the prevalence and median abundance of the identified VFCs exhibited a general trend of the more prevalent VFCs also being more abundant in gut metagenomes (**Fig. 1D**). Strikingly, the most prevalent VFCs in our cohort do not correspond to well-known families such as crAssphages^22–24^ and Gubaphage/*Flandersviridae*^17,32^, with all 12 highly prevalent VFCs (>50% prevalence) containing no references with ICTV-ratified family assignments, and only the most prevalent VFC (VFC 1; VFCs are numbered based on prevalence in our cohort) being related to a lineage that has been described in a previous study^26,50^ (*Heliusvirales*; **Fig. 1D**). The newly identified VFCs typically include members of known vOTUs but also include a substantial fraction of vOTU diversity that is novel (>47% on average; **Fig. 1D**), highlighting the value of a long-read assembled vOTU catalog in their construction. Functional annotation of key viral structural proteins (portal, large terminase, and major capsid protein; **Supplementary Fig. 7**; **Supplementary Data 4**), and analysis of phylogenetic diversity among the most prevalent VFCs, confirmed that they exhibit robust clustering patterns similar to known viral clades (**Supplementary Fig. 8**). Visualization of the phylogenetic tree for the portal protein identified viral clades that closely mirror VFC clusters constructed here as well as known viral families (**Fig. 1E**), further supporting the robustness of this approach to delineate candidate viral families. Similar clades were seen also for the large terminase and major capsid proteins, with some evidence for subfamily-level divergence and potential horizontal gene transfer events (**Supplementary Fig. 9**). Overall, these observations suggest that highly prevalent viral clades in our cohort are not currently well-represented by known taxa, many of which have been defined based on isolate genomes and short-read metagenomic assemblies.

Working with the initial hypothesis that the highly prevalent VFCs could represent population-specific viral diversity, we sought to assess their prevalence based on global metagenomic datasets (n=3,011, restricted to deeply sequenced gut metagenomes i.e. >50M reads; **Methods**; **Supplementary Data 5**). Surprisingly, several of our VFCs were found to be among the most highly prevalent globally as well (**Fig. 1F**), with largely similar patterns seen across different continents (**Supplementary Fig. 10**). In particular, VFC 3 was consistently found to be the most prevalent globally, with 6 out of 12 newly identified VFCs from this study being among the globally prevalent families (prevalence >20%; **Fig. 1F**). Intriguingly, while for most VFCs prevalence was driven by the cumulative contribution of many vOTUs, VFC 3 is unique in that it has a few highly cosmopolitan vOTUs (**Supplementary Fig. 11**). Consistent with prior reports, well-known phage families such as crAssphages and Gubaphage/*Flandersviridae* are indeed among the most abundant in global datasets (though they are biased towards western populations; **Supplementary Fig. 12**), while the prevalent VFCs described here are more abundant in our population (**Fig. 1D**), highlighting that population-specific studies can indeed uncover gut phage families that may be more important in specific regions of the world. Overall, these results highlight the surprising discovery of globally highly prevalent phage family-level clusters based on a population-specific long-read gut metagenomics dataset, emphasizing the need for more such studies in populations around the world.

### Broad host ranges and Firmicutes hosts characterize many highly prevalent gut phage families

To characterize potential hosts for the phage families identified here, a combination of three different approaches was leveraged to increase sensitivity while maintaining specificity (**Fig. 2A**). In particular, spacer matching and prophage analysis are commonly used bioinformatic approaches that can benefit from the contiguity and quality of long-read**–**based MAGs^37^ and vOTUs. Correspondingly, clear advantages were seen for metagenomic assemblies augmented with long reads in terms of identification of CRISPR cassettes and prophage hosts^36^ (167% increase in cassettes detected and >10× increase in prophage sequence context assembled; **Supplementary Fig. 13**). In addition, extensive proximity-ligation sequencing data (Hi-C, n=108) was used to provide complementary experimental evidence for identifying phage-host linkages that may not be captured by bioinformatic techniques alone^51,52^ (**Fig. 2A**). Based on a model that accounts for noise from extracellular DNA, highly significant phage-host linkages were identified from this data (**Supplementary Fig. 14**), which in combination with spacer matching and prophage analysis enabled host associations being mapped for a majority of vOTUs (>58%) and nearly all VFCs (>96%). Of note, Hi-C data provided host information for the largest number of vOTUs (>14,000 versus <8,000 for spacer matching and prophage analysis), with >5,000 vOTUs for which only Hi-C data identified host linkages (**Supplementary Fig. 15**; **Supplementary Data 6)**.

**Figure 2.**
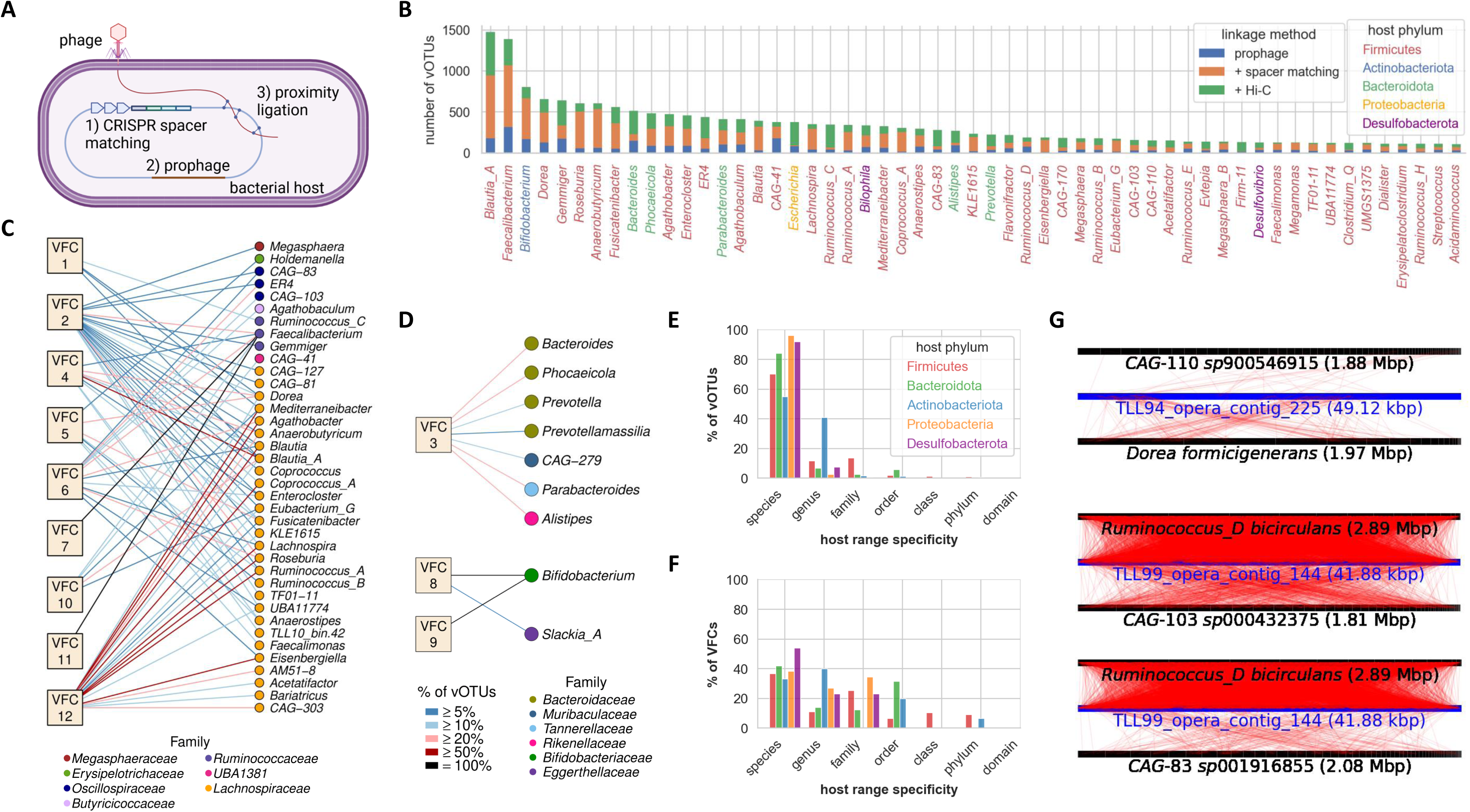
Comprehensive host association analysis reveals Firmicutes hosts and broad host range as common features of highly prevalent gut phage families. (A) Schematic of approaches used to predict hosts of the vOTUs (created with BioRender.com). Hosts were predicted for each viral sequence and aggregated per vOTU (**Methods**). (B) Host genus distribution coloured by phylum and showing the number of vOTUs linked when prophage linkage, spacer matching, and Hi-C information are sequentially added. (C-D) Host genera linked to members of the 12 most prevalent VFCs, with Firmicutes hosts (C), Bacteroidota and Actinobacteriota hosts (D) shown separately. VFC-genus linkages are color-coded by the percentage of vOTUs in the VFC linking to that genus. Hosts are colored by family. (E) Host range distribution of vOTUs stratified by host phylum. (F) Host range distribution of VFCs stratified by host phylum. Firmicutes-infecting VFCs have significantly broader host ranges than the others (p=0.002, one-sided Mann-Whitney U test). (G) Hi-C evidence for broad host ranges of individual viral *genomes*, for two different viral contigs belonging to VFC 1 and their host MAGs. Inter-MAG linkages do not exceed 10, indicating little cross-contamination between MAGs and suggesting genuine inter-genus interactions.

The top 10 most common gut microbial genera that were identified as hosts belonged to the phyla Firmicutes (*Blautia_A*, *Faecalibacterium*, *Dorea*, *Anaerobutyricum*, *Gemmiger*, *Roseburia*, *Fusicatenibacter*), Actinobacteria (*Bifidobacterium*) and Bacteroidota (*Bacteroides*, *Phocaeicola*; **Fig. 2B**), consistent with the relative prevalence of these genera in the cohort and the number of MAGs assembled (**Supplementary Fig. 16**). In comparison, vOTU association counts in a genus were less correlated with the median abundance of the genus or the abundance of associated vOTUs (**Supplementary Fig. 16**), indicating that host prevalence plays a stronger role in determining the presence of vOTUs. Overall, evidence for host associations across genera were provided by all three approaches used, though some genera had biases towards spacer matching (e.g. *Anaerobutyricum*, *Lachnospira*) or Hi-C data (e.g. *Escherichia*, *Rumincoccus_C*), likely as a reflection of the uneven prevalence of CRISPR-Cas systems across taxa^53^ and the inherent biases of these methods in identifying historical versus active infections^54^ (**Fig. 2B**).

Most vOTUs (>84%) were associated with hosts from a single genus, though evidence from spacer matching and Hi-C data supported broader host ranges in some cases (15% of vOTUs infect >1 genera; **Supplementary Fig. 17**). As expected, VFC host ranges were broader than those of their constituent vOTUs (**Supplementary Fig. 17**), although each has a predominant host phylum (**Supplementary Fig. 18**). Among the most prevalent VFCs, 9 out of 12 (VFCs 1, 2, 4, 5, 6, 7, 10, 11 and 12) broadly infect members of the Firmicutes phylum, especially members of the *Lachnospiraceae* family (**Fig. 2C**), and we therefore refer to them as *GuFi* (Gut Firmicutes) phages. The wide host range of VFC 1 (*Heliusvirales*) has been reported previously^26^, with VFCs 2, 4, 5, 6, 10 and 12 having similarly broad host ranges, while VFCs 7 and 11 almost exclusively infect *Faecalibacterium* species. In contrast, VFC 3 infects Bacteroidota hosts, while VFCs 8 and 9 infect Actinobacteriota, most notably *Bifidobacterium* species (**Fig. 2D**). These results suggest that host range breadth and the prevalence of the host both contribute to the prevalence of a VFC. While the majority of vOTUs have species or genus-specific host ranges (**Fig. 2E**, **Supplementary Fig. 17A**), Firmicutes-infecting VFCs have significantly broader host ranges than VFCs infecting other phyla (p=0.002, one-sided Mann-Whitney U test; **Fig. 2F**). In addition, the 12 most prevalent VFCs in this study have >4× greater host diversity relative to other VFCs (family-level entropy=1.3 versus 0.3; **Methods**). Having broader host range in a VFC was also found to be associated with higher prevalence in the population (**Supplementary Fig. 19**). Additionally, we found evidence of individual viral genomes having broad host ranges, including a VFC 1 phage that has Hi-C linkages to 2 MAGs from different orders (upper panel, **Fig. 2G**), and another to 3 MAGs from 2 different families (middle and lower panel, **Fig. 2G**). The linkages are broadly distributed over both the viral genome and each MAG, and 10 or fewer linkages between the hosts targeted indicate little cross-contamination between the MAGs. These results are in line with recent observations^52^ and suggest that phages can interact with a much broader range of hosts than is commonly appreciated.

### Highly prevalent VFCs actively replicate in the human gut

Phages are often either obligately lytic (virulent), replicating and lysing their hosts, or have the ability to integrate into the host genome as dormant prophages (temperate), with the potential to excise and re-enter the lytic cycle or eventually degrade into a defective state^55^. Among the most prevalent VFCs, the Bacteroidota-infecting VFC 3 has the lowest incidence of vOTUs identified as being temperate based on gene annotations (18%), whereas the Firmicutes and Actinobacteriota VFCs frequently have temperate vOTUs identified (61-94%; **Methods**). For prophages, it is often unclear to what extent they actively replicate in response to environmental triggers^56–58^, particularly in the *in vivo* state in the human gut microbiome. To address this, we combined multiple complementary analysis approaches, as well as virus-like particle (VLP) enrichment and sequencing of samples (n=64), to systematically identify actively replicating phages (**Fig. 3A**; **Supplementary Fig. 20**; **Methods**). This included identification of vOTUs with high mean read coverage relative to their linked MAGs in (i) bulk metagenomics^59,60^ and (ii) VLP sequencing data, as well as vOTUs with outlier coverage relative to non-viral contigs when assessed in (iii) VLP sequencing data or (iv) as reads per kilobase million (RPKM) ratios between VLP and bulk metagenomic sequencing data (**Fig. 3A**; **Methods**; **Supplementary Data 7-8**). This integrated approach identified >6,000 vOTUs (out of 30,510, >20%) with replication signatures in at least one sample, and >100 vOTUs/sample on average with evidence for replication. While the different approaches had significant overlap in the vOTUs identified as replicating, they also exhibited clear complementary strengths (**Supplementary Fig. 21-22**).

**Figure 3.**
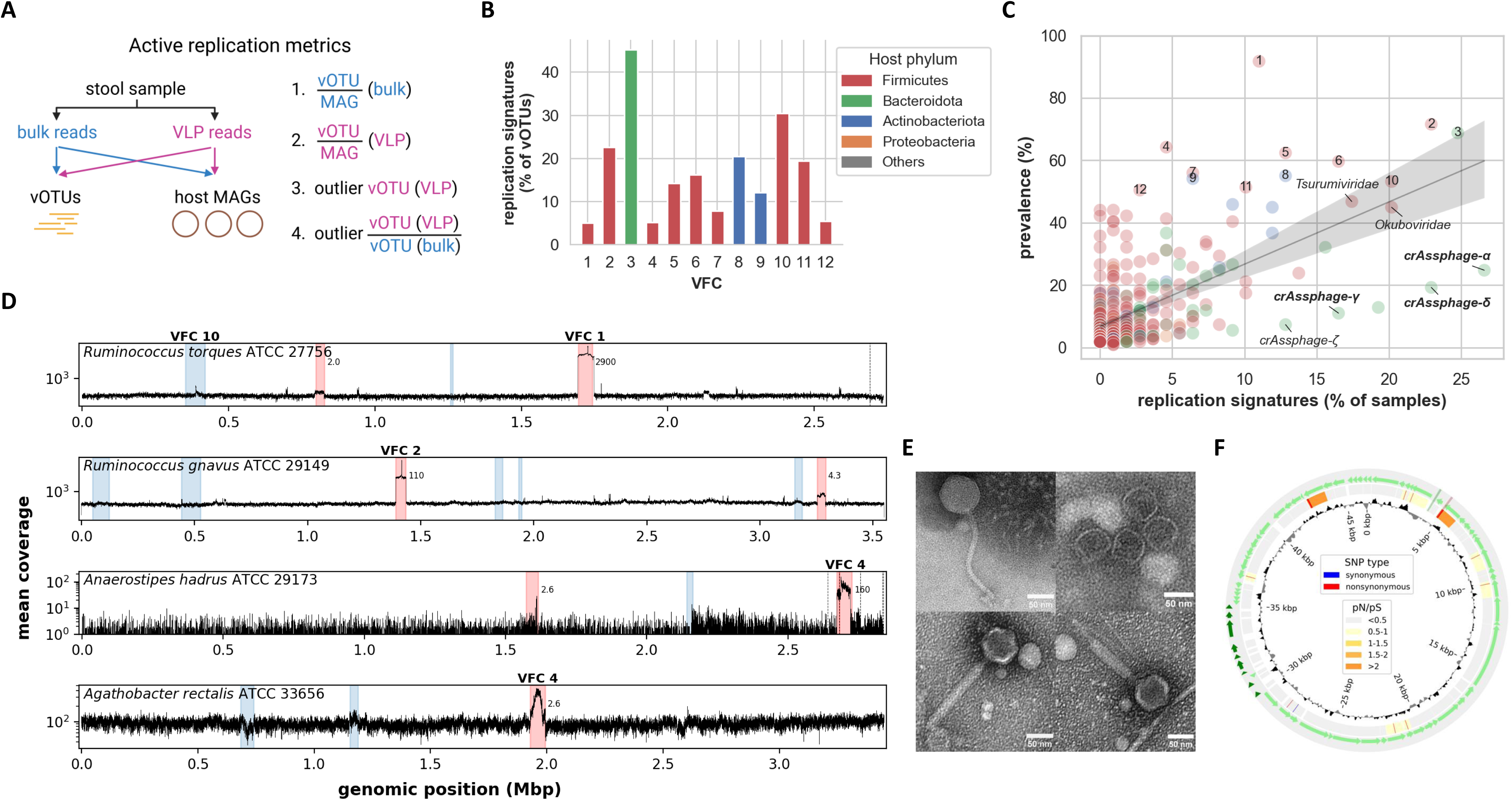
Highly prevalent gut phage families actively replicate in the human gut. (A) Schematic of approaches used to identify signatures of actively replicating vOTUs based on read coverage analysis. (B) Distribution of vOTUs with replication signatures in at least 1 sample in VFCs 1-12. (C) Distribution of VFC prevalence versus percentage of samples in which the VFC contains at least one actively replicating vOTU. VFCs with >50% prevalence are numbered (1-12). Reference families replicating in >10% of samples are labelled, with ICTV-ratified families in bold. VFCs 2 and 3 were detected to be actively replicating in the most samples, together with *Crassvirales* families. The VFCs are color-coded by host phylum. (D) VFC 1-4 prophages induce *in vitro*. Bacteria were cultured with the addition of mitomycin C, VLP DNA was sequenced, and reads mapped back to the genome assemblies (**Methods**). Prophage regions with mean coverage >2× of the host are highlighted in red and the mean ratios are indicated; other predicted prophages are highlighted in blue. Vertical dotted lines indicate contig boundaries. *Ruminococcus torques* VFC 1 phage, *Ruminococcus gnavus* VFC 2 phage, *Anaerostipes hadrus* VFC 4 phage, and *Agathobacter rectalis* VFC 4 phage were all detected to be actively replicating. (E) Transmission electron microscopy images of *R. torques* VFC 1 phage (upper left), *R. gnavus* VFC 2 phage (upper right), and *A. hadrus* VFC 4 phage with non-contracted (lower left) and contracted tails (lower right). Scale bars are shown on the lower right of each image. (F) Circos plot of a VFC 2 vOTU with active DGRs. The outer track shows genes (annotations and PHROG categories in **Supplementary** Fig. 29A), the middle track shows synonymous (blue) and nonsynonymous (red) mutations and pN/pS ratio of genes (darker shades of orange indicate higher pN/pS; **Methods**), and the inner track shows coverage skew. The DGR template (green) and variable (red) regions are highlighted spanning the outer and middle tracks.

Focusing on the most prevalent VFCs, 45% of VFC 3 vOTUs and 5-30% of the others were detected to be replicating in at least one sample (**Fig. 3B**), with evidence for replication typically being detected using multiple methods (**Supplementary Fig. 22**). While overall there appears to be a correlation between VFC prevalence and the fraction of samples with replication signatures, several of the top VFCs (e.g. 1, 4, 7, 9 and 12) are more prevalent than expected, while several crAssphages families are less prevalent (**Fig. 3C**). Of note, most of the top VFCs (8 out of 12) exhibit replication signatures in a similar number of subjects as crAssphages families (10-30%; **Fig. 3C**), emphasizing that despite being often annotated as being temperate, many of the GuFi VFCs can frequently impact their hosts through active replication.

### Members of the most prevalent VFCs readily induce from their hosts *in vitro*

To experimentally confirm that prophages of highly prevalent VFCs induce from their hosts, we sourced publicly available Firmicutes strains carrying members of these families as prophages: *Ruminococcus torques* ATCC 27756 (VFC 1), *Ruminococcus gnavus* ATCC 29149 (VFC 2), *Anaerostipes hadrus* ATCC 29173 (VFC 4), and *Agathobacter rectalis* ATCC 33656 (VFC 4; **Supplementary Table 1**; **Methods**). The isolates were then cultured with and without addition of an inducing agent (mitomycin C) and induced prophages were detected using a range of methods (**Supplementary Fig. 23A**). PCR primers were designed to detect each internal phage region and the circularized phage genome^36^ (junction of circularization or JoC; **Supplementary Fig. 23B**; **Supplementary Table 3**). The presence of the JoC band for all 4 phages in genomic DNA indicated that all of them successfully excise from the host genome and circularize. Furthermore, the presence of both the phage region and JoC bands in DNase-treated VLP DNA, together with the lack of bacterial contamination, suggests that all form complete phage particles as well (**Supplementary Fig. 24**). Sequencing analysis of VLP filtrates showed clear evidence for induction of all phages, with high prophage mean coverages relative to the chromosomal regions for all except for *A. rectalis* (>100×; **Fig. 3D**). In fact, the prophages were found to readily induce even without the addition of mitomycin C, indicating that this could be happening *in vivo* as well (**Supplementary Fig. 25**). Non-uniformities in read coverage within the phage regions were observed (**Supplementary Fig. 26A**), which were consistent with a cos (3’) packaging strategy for the *R. torques* VFC 1 phage and a headful (pac) packaging strategy for the *R. gnavus* VFC 2 phage, respectively (**Supplementary Fig. 26B**).

Further scale-up of cultures and concentration of phage preparations was done to enable viewing under a transmission electron microscope (TEM; **Methods**). We observed complete phage particles (**Fig. 3E**, **Supplementary Fig. 27**) of the *R. torques* induced VFC 1 phage exhibiting a long, flexible tail morphology and a characteristic icosahedral head (*Siphoviridae*; **Fig. 3E**, upper left). In contrast, the *A. hadrus* induced VFC 4 phage exhibits both contracted and non-contracted tail forms consistent with a *Myoviridae* morphology (**Fig. 3E**, lower left and lower right). Interestingly, capsid-like structures were observed for the *R. gnavus* induced VFC 2 phage, but these appear to be empty in the TEM images (**Fig. 3E**, upper right). No particles of the *A. rectalis* induced VFC 4 phage could be viewed, potentially due to the low induction rate in the conditions tested. Nevertheless, these images represent the first visual evidence for phage particles of some of the most prevalent phage families in the human gut microbiome.

### Distinguishing genomic features for highly prevalent VFCs

We annotated genomic features for the top VFCs in the search for attributes that may explain their wide distribution in the human gut. Several VFCs frequently carry diversity generating retroelements (DGRs), especially VFCs 2, 3, 4, 6, 10 and 12 having 15-72% incidence and being outliers compared to other VFCs (**Supplementary Fig. 28A**; **Supplementary Data 9**). To check to what extent the DGRs are functional, we identified single nucleotide variants (SNVs) for vOTUs within a metagenome, and computed the relative rates of synonymous versus nonsynonymous SNVs within targeted and non-targeted genes (pN/pS ratio; **Methods**; **Supplementary Data 10**). Among the most prevalent VFCs, we found a statistically higher pN/pS ratio for genes containing DGR variable regions relative to other genes (median 0.9 vs 0.3; two-sided Mann-Whitney U test p-value<10^-6^; **Supplementary Fig. 28B**), suggesting that the DGRs tend to be active and promote high rates of diversifying mutations in their target genes. The most common annotations of target genes include “head decoration protein” and “tail fibre protein”, indicating roles in both immune evasion and broadening host range for DGRs (**Supplementary Fig. 28C**). For example, in a DGR-carrying VFC 2 vOTU, the annotated DGR and another gene exhibit high pN/pS ratios with 10-11 nonsynonymous mutations within the final 100 bp of the genes, while there are a total of 9 SNVs elsewhere in its 46 kbp genome (**Fig. 3F**, **Supplementary Fig. 29A**). This vOTU has Hi-C linkages to two host *Gemmiger* species, highlighting its ability to infect distinct species. As another example, a VFC 4 vOTU was found to have 9 nonsynonymous mutations in the DGR variable region but just 5 elsewhere in the >60 kbp replicating region, along with Hi-C linkages to hosts from two different genera (*Mediterraneibacter faecis* and *Fusicatenibacter saccharivorans*; **Supplementary Fig. 29B**). These examples illustrate how DGRs may help enhance the ability of members of these prevalent families to disseminate to new hosts in the human gut.

The top VFCs were also found to carry a diverse array of systems that combat bacterial antiphage defences such as restriction-modification and CRISPR-Cas systems (VFC 3 is also an outlier in this; **Supplementary Fig. 30A**). Intriguingly, diverse antiphage defence systems were also detected among the vOTUs, suggesting functions in inter-phage warfare^61–64^ (VFC 1 is an outlier in this; **Supplementary Fig. 30B**). The top VFCs also carry a range of auxiliary metabolic genes (AMGs) that reprogram the metabolic capacities of their hosts (**Supplementary Fig. 31A**). VFC 1 stands out in its incidence of restriction-modification systems and methyltransferases. In fact, the protein cluster containing methyltransferase annotations is carried by >97% of VFC 1 vOTUs (**Supplementary Data 11**). We hypothesize that this unique feature is associated with the curious observation that <4% of VFC 1 vOTUs have CRISPR spacers matching them, in contrast to other VFCs with similar host association rates (**Supplementary Fig. 31B**). While a high incidence of methyltransferases has been observed previously^26^, a potential DNA-modification-based function for evading CRISPR-Cas immunity for this widespread viral clade has not been suggested before. We identified a VFC 1 vOTU carrying a restriction-modification system that was also linked by Hi-C to two different species (**Supplementary Fig. 32**).

Combining genetic features (e.g. DGR prevalence, anti-defence system prevalence, etc.) with the other attributes identified in this study as potentially explaining VFC prevalence (e.g. host diversity, Firmicutes hosts, etc.; **Supplementary Fig. 33**), we constructed a generalized linear model (GLM) as well as a random forest regression (RFR) model for VFC prevalence (**Methods**). Both models consistently highlighted host (and VFC) abundance and diversity (family and order-level entropy) as the key attributes explaining the prevalence of VFCs (variance explained=48% for GLM and 21% for RFR). In addition, other features such as having Firmicutes hosts, the percentage of samples with a replication signature, and frequency of DGR, anti-defence, AMG and phage lifestyle annotations also contributed to the models, indicating that these features may also impact the observed prevalence of a VFC. Overall, this provides evidence that the high prevalence of GuFi phages in the human gut microbiome may be explained by many factors including host diversity, phage lifestyle, and genetic attributes that enable survival in a highly contested environment.

## Discussion

The present study demonstrates that long-read metagenomics provides a transformative advantage for the discovery and characterization of DNA phage diversity in the human gut. Although tailed phage genomes are typically small as compared to cellular organisms (10-150kbp)^65^, several intrinsic features—including their low abundance, high repeat content, and substantial intra-host heterogeneity—can contribute to the fragmentation and loss of viral sequences in short-read assemblies^66–68^. By leveraging hybrid assemblies in which long reads augment accurate short-read contigs, we have shown that long-read data markedly improves viral genome contiguity and completeness (**Fig. 1B**), thereby enabling more reliable vOTU recovery (**Supplementary Fig. 3**) and robust construction of viral family-level clusters (**Supplementary Fig. 5-8**) across 109 deeply sequenced individuals in a multi-ethnic Asian cohort. Notably, 63% of our reconstructed vOTUs were absent from eleven major public gut virome datasets (**Fig. 1C**), highlighting persistent underrepresentation of gut viral diversity in short-read–derived catalogs, in addition to the high individual-level specificity of gut viromes^44,45^. As long-read sequencing technologies continue to decrease in cost and increase in throughput, deep long-read metagenomic profiling of hundreds of samples may offer greater scientific value than short-read profiling of thousands, especially given that participant recruitment—rather than sequencing—is often the major bottleneck in population microbiome studies.

A striking outcome of this work is the discovery of several highly prevalent and deeply branching viral lineages that have been largely overlooked in prior surveys (**Fig. 1D–E**). While genomic analyses have long suggested an abundance of prophages in Firmicutes genomes^17,69^, the prevailing emphasis in the literature has centered on phages infecting Bacteroidota, in part because these lineages are more easily recovered from viral-like particle–enriched datasets^23,45,70^. Our results reveal that this historical imbalance has obscured the global relevance of several Firmicutes-infecting phage families that we refer to as GuFi phages. Notably, our data rediscovers the *Heliusvirales* lineage^26,50^ (VFC 1) and uncovers additional previously uncharacterized clades including VFC 2 (proposed name *Astravirales*) and VFC 4 (proposed name *Suryavirales*) that are globally widespread. Conversely, we also report a newly identified Bacteroidota-infecting lineage (VFC 3; proposed name *Tarakavirales*) that appears to be one of the most prevalent gut phage clades worldwide (>80%) with highly cosmopolitan vOTUs (**Supplementary Fig. 11**), suggesting that both population-specific and globally distributed phage clades remain to be systematically characterized. We note the technical difficulties inherent in defining novel viral clades beyond the genus level, that exist even despite improved genome construction enabled by long-read sequencing, due to complicated phylogenetic relationships and genetic mosaicism between numerous phage clades^27,28^. While we have striven to construct family-level clusters with good agreement with known, reference families (**Supplementary Fig. 5**), we find that several of the most prevalent VFCs have high phylogenetic diversity that belies calling them “families” (**Supplementary Fig. 8-9**). Thus, we believe VFCs 2-4 are more appropriately classified at the order level, similar to *Heliusvirales*^26^, and have named them as such (ending with “-virales”).

A defining feature of several GuFi phage families is the breadth of their predicted host range, supported by an integrated annotation framework combining CRISPR spacer matching, prophage–host genomic adjacency, and Hi-C–based linkage (**Fig. 2B–D**). Layering additional host-prediction signals markedly increases assignment confidence and reveals widespread host breadth among these lineages (**Supplementary Fig. 15, 17**). While phage host ranges are often described as narrow in traditional laboratory settings^71,72^, Hi-C data provided direct evidence that individual GuFi phage genomes establish linkages to multiple bacterial genera^52^ (**Fig. 2G**), where low inter-MAG Hi-C linkages (<10) indicate little cross-contamination and support genuine multi-genus host interactions. At the VFC level, Firmicutes-infecting families have vOTUs that exhibit significantly broader host ranges than Bacteroidota- or Actinobacteriota-infecting families (p=0.002; **Fig. 2E-F**), suggesting that broad tropism may facilitate persistence and dissemination in the gut ecosystem, particularly for temperate phages that must encounter susceptible hosts amid fluctuating microbial communities.

Multiple lines of evidence indicate that prevalent GuFi phage clades are actively replicating *in vivo*. VLP sequencing identified replication signatures for vOTUs belonging to prevalent GuFi VFCs in a substantial fraction of samples (**Fig. 3B–C**; **Supplementary Fig. 20–22**). Several GuFi VFCs display replication frequencies comparable to those of lytic crAssphage families, despite encoding canonical markers of lysogeny and having distinct host ranges and genomic architectures. Of note, lysogenized phages may be more challenging to detect replication signatures for as induced members are not as abundant as strictly lytic phages, so our analysis likely represents an underestimate. Controlled induction experiments further demonstrate that prophages from VFCs 1, 2 and 4 readily enter the lytic cycle *in vitro*—even without classical stressors^73^—producing abundant VLPs (**Fig. 3D**; **Supplementary Fig. 24-25**). Similar induction rates *in vivo* likely confer ecological advantages to the phage and contribute to their high prevalence. Transmission electron microscopy confirms that these induced particles possess typical *Caudoviricetes* morphologies, including contracted-tail states for some members (**Fig. 3E**; **Supplementary Fig. 27**). Together, these results demonstrate that globally prevalent Firmicutes phage families are not dormant genetic elements but dynamic and frequently replicating members of the gut ecosystem.

Genomic analyses further reveal that GuFi lineages encode a rich suite of adaptive features that likely contribute to their ecological success. Diversity-generating retroelements (DGRs)—previously shown to be common and active in gut phages^74,75^—are enriched across several VFCs, frequently targeting tail fiber, head decoration, or receptor-binding proteins (**Fig. 3F**; **Supplementary Fig. 28–29**). Elevated pN/pS ratios at DGR-associated loci indicate ongoing diversification that may modulate host specificity or immune evasion^74,76^. Many VFCs also encode anti-defence elements, including predicted CRISPR evasion or abortive infection components (**Supplementary Fig. 30**), and auxiliary metabolic genes, including predicted methyltransferases in VFC 1 that may protect against host restriction systems and potentially even CRISPR-Cas systems^77–79^ (**Supplementary Fig. 31-32**). These results show that prevalent families have evolved multilayered strategies to overcome host immunity and persist in densely competitive gut ecosystems.

Together, these findings reshape current models of gut virome community structure derived from short-read-centric surveys. Long-read metagenomics enables more comprehensive reconstruction of full-length viral genomes, revealing globally dominant phage clades—many infecting Firmicutes—that have been systematically overlooked in previous studies. The broad host ranges, active replication, and extensive anti-defense and diversification mechanisms encoded by GuFi phage families position them as key modulators of gut microbial ecology. Expanding long-read metagenomic surveys, especially in understudied populations, will be essential for resolving the global landscape of human gut viral diversity and elucidating the roles of dominant phage lineages in microbiome stability, microbial competition, and human health.

## Methods

### Subject recruitment

Subjects for this study were recruited based on recall from a community-based multi-ethnic prospective cohort^80^ as described previously^37^ (n=109). They comprise 65 males and 44 females, 48 to 76 years old at the time of sample collection, with no pre-existing major health conditions (cardiovascular disease, mental illness, diabetes, stroke, renal failure, hypertension, or cancer) based on self-reporting^80^. Informed consent was obtained from all participants. All associated protocols for this study were approved by the National University of Singapore Institutional Review Board (IRB reference number H-17-026).

### Sample collection and DNA extraction

Faecal samples were collected from the subjects as described previously^37^. Briefly, samples were collected using the BioCollector^TM^ kit (The BioCollective, Colorado, USA), brought to the Temasek Life Sciences laboratory and stored in an anaerobic chamber (atmosphere of N_2_ (75%), CO_2_ (20%), and H_2_ (5%)) within the same day. Faecal samples were homogenized and subsamples transferred into sterile 2 mL centrifuge tubes. Genomic DNA was extracted from faecal material (0.25g wet weight) using the QIAamp Power Faecal Pro DNA kit (QIAGEN GmbH, Cat. No. 51804) and quantified using the Qubit dsDNA BR Assay Kit (Thermo Fisher Scientific, Cat. No. Q32853).

### Illumina and ONT library preparation and sequencing

Illumina and ONT metagenomic libraries were prepared as described previously^37^. Illumina paired-end sequencing (2×151bp reads) was performed on the HiSeq4K platform, with a minimum and median depth per sample of 2.4Gbp and 9.5Gbp, respectively. Following ONT library preparation, single-plex samples were sequenced on either a MinION or GridION machine with either FLO-MIN106D or MIN106 revD flowcells. Multiplex samples were sequenced on a PromethION machine with FLO-PRO002 flowcells. Raw ONT reads were basecalled using the latest version of the basecaller available at the point of sequencing (Guppy v3.0.4 to v3.2.6). Basecalled reads were demultiplexed using qcat (https://github.com/nanoporetech/qcat) v1.1.0. The minimum and median sequencing depth for ONT reads per sample were 1.8Gbp and 5.7Gbp, respectively.

### Metagenomic assembly, binning, and taxonomic classification

Short-read metagenomic assemblies were generated from Illumina reads using MEGAHIT^81^ v1.0.4-beta. Hybrid metagenomic assemblies were generated from Illumina and ONT reads using OPERA-MS^38^ v0.9.0, where the MEGAHIT assemblies provided contigs that were further scaffolded and extended. Assemblies were binned to form metagenome-assembled genomes (MAGs) using MetaBAT2^82^ v2.12.1. MAGs were evaluated for completeness and contamination using CheckM^83^ v1.0.4, and those with completeness >50% and contamination <10% (n=4,497, medium-quality and above by MIMAG standards^84^) were retained for further analysis. MAGs were compared with GTDB^85^ release 95 using the ‘ani_rep’ command from GTDB-Tk^86^ v1.4.1, which computes pairwise Mash^87^ distances between all query MAGs and GTDB genomes. Mash distance ≤0.05 was used for taxonomic classification of each MAG.

### Viral identification and dereplication

Viral sequences were identified from the metagenomic assemblies using 2 approaches: (1) VirSorter2^39^ v2.2.3 (options ‘--keep-original-seq’) followed by CheckV^40^ v0.8.1, retaining sequences with ‘viral_genes’ ≥ ‘host_genes’; (2) geNomad^41^ v1.6.1 (options ‘--enable-score-calibration’) retaining sequences with ‘fdr’ <0.01. Sequences identified via either approach and of length ≥5 kbp were combined, and dereplicated using CD-HIT^88^ v4.8.1 at 99% global identity (options ‘-c 0.99’) followed by 95% identity over 85% of the shorter sequence (options ‘-c 0.95 - G 0 -aS 0.85’) following MIUViG standards^42^. 30,510 viral operational taxonomic units (vOTUs) were recovered after dereplication.

### Genome completeness and lifestyle prediction

vOTU completeness was estimated using CheckV^40^ v1.0.1. To identify circular sequences, Illumina and ONT reads were assembled using Unicycler^89^ v0.5.0 (with the OPERA-MS assembly provided to the ‘--existing_long_read_assembly’ option). Unicycler contigs with the “circular=true” flag were compiled and aligned to the viral sequences using blastn^90^ v2.14.0 (options ‘-evalue 1e-12 -perc_identity 95 -max_target_seqs 10000’). Alignments between each pair of sequences were merged using ‘anicalc.py’ from the CheckV package^40^, and circular vOTUs were identified if alignments cover ≥85% of both the query and target. Viral lifestyles (virulent or temperate) were predicted using BACPHLIP^91^ v0.9.6.

### Long and short-read viral recovery comparison

An identical viral identification pipeline was run on the short-read metagenomic assemblies constructed using MEGAHIT v1.0.4-beta to generate viral sequences and vOTUs. For viral recovery comparison, long and short-read viral sequences were combined, and dereplicated using CD-HIT v4.8.1 at 99% global identity (options ‘-c 0.99’) followed by 95% identity over 85% of the shorter sequence (options ‘-c 0.95 -G 0 -aS 0.85’). This procedure produced 34,860 jointly dereplicated clusters containing only short-read sequences, long-read sequences, or both.

### Novelty analysis

vOTUs were combined with the GVD^16^, GPD^17^, MGV^18^, CHVD^19^, IMG/VR v4^20^, CHGV^29^, DEVoC^30^, and RefSeq Viral^31^ databases, and the datasets of Benler et al.^32^, Nishijima et al.^25^, and Shah et al.^33^, and an all-versus-all sequence alignment was performed using blastn^90^ v2.14.0 (options ‘-evalue 1e-12 -perc_identity 95 -max_target_seqs 10000’). Alignments between each pair of sequences were merged using ‘anicalc.py’ from the CheckV package^40^. A database sequence was considered a vOTU match if an alignment covered ≥85% of the vOTU. To estimate the number of vOTUs present in the 11 public datasets, the viral sequences were combined and dereplicated using Vclust^92^ v1.3.1 in a three-step process (‘vclust prefilter’ with options ‘--min-ident 0.95 --batch-size 2000000 --kmers-fraction 0.2 --min-kmers 4’; ‘vclust align’ with options ‘--out-ani 0.95 --out-qcov 0.85’; and ‘vclust cluster’ with options ‘--algorithm cd-hit --metric ani --ani 0.95 --qcov 0.85 --out-repr’).

### Viral gene identification and functional annotation

Gene calling was performed using prodigal-gv^41^ v2.11.0, a fork of Prodigal^93^ incorporating models for alternative genetic codes. Functional annotation was performed using the ‘genomad annotate’ command^41^, which aligns proteins to a set of virus, plasmid, and chromosome-specific markers annotated with PFam, COG, TIGRFAM, and KEGG orthology accessions. Proteins were separately annotated using pharokka^94^ v1.7.4, which aligns proteins to the PHROG database^95^.

### Viral family-level cluster construction

Viral sequences from Yutin et al.^24^, Benler et al.^32^, Nishijima et al.^25^, Shah et al.^33^, and de Jonge et al.^26^, with proposed taxonomic assignments for novel gut phage lineages, were combined with the ICTV Master Species List 39^47^ (“host source” of Bacteria or Archaea only, segmented genomes (n=13) and gene transfer agents (n=7) removed), and dereplicated at 95% identity over 85% coverage of the shorter sequence to generate a set of reference vOTUs. Genome completeness was estimated using CheckV, and genes were identified using prodigal-gv as above. Reference vOTUs with ≥50% completeness (n=16,860) were combined with ≥50% complete SPMP vOTUs (n=7,484), and an all-versus-all protein sequence alignment was performed using DIAMOND^96^ v2.1.8 (options ‘--more-sensitive -e 1e-5 --max-target-seqs 10000’). Proteins were clustered on the basis of alignment identity × alignment length to form protein clusters (PCs) using MCL^97^ v22.282 (options ‘-I 2’). A total of 85,419 non-singleton PCs were obtained. The number of PCs shared between every pair of sequences was then computed.

High-quality (≥90% complete) and complete reference and SPMP vOTUs were used to form viral clusters. UPGMA hierarchical clustering was performed using 100 minus the percentage of shared distinct PCs as the distance metric. Inspection of reference families led to a cutoff of 20% shared PCs to define family-level clusters (**Supplementary Fig. 5**; **Supplementary Data 2**). Other vOTUs (50-90% complete) were recruited into the VFCs if their mean shared PCs with high-quality or complete members was at least 30%, a threshold chosen to minimize ambiguous assignments to >1 VFC. This resulted in 351 non-singleton viral family-level clusters (VFCs) containing at least two SPMP vOTUs.

### Phylogenetic analysis

Phylogenetic analyses were performed on the large terminase (LT), portal protein, and major capsid protein (MCP) of highly prevalent VFCs and selected reference families. For each VFC, protein clusters (PCs) were manually inspected and those containing sequences annotated as LT, portal, or MCP by geNomad and/or pharokka were assigned those functions. Length thresholds of 900bp, 900bp, and 700bp for LT, portal, and MCP, respectively, were implemented. Up to 3 high-quality (≥90% complete) or complete vOTUs or reference genomes were randomly selected per subclade sharing 40% PCs (‘genus-level’), picking sequences carrying all annotations if possible. Multiple sequence alignments were constructed using MAFFT^98^ v7.520 (‘mafft-linsi’) and trimmed using trimAl^99^ v1.4.1 (options ‘-automated1’). Phylogenetic trees were built using FastTree^100^ v2.1.11 and visualized on the iTOL^101^ webserver. The phylogenetic dispersion of a clade was computed as the maximum pairwise distance of members in that subclade.

### Prevalence and abundance estimation

For assessing prevalence in the SPMP cohort, short reads from each sample were mapped to all vOTUs using bwa-mem^102^ v0.7.17. Reads were filtered to retain properly paired mappings at ≥95% identity over ≥80% of the read length. A vOTU was considered present in a sample if reads cover ≥70% of its length. To estimate relative abundances, the mean coverage of each vOTU was computed excluding the top and bottom 10% of base coverage values, and the relative abundance was calculated as the mean divided by the sum of mean coverages for all vOTUs detected in the sample. A VFC was considered present in a sample if at least one vOTU belonging to that VFC was detected. The relative abundance of a VFC was taken to be the sum of relative abundances of its constituent vOTUs. The VFCs are numbered according to prevalence in the SPMP cohort (VFC 1 is the most prevalent, etc.).

For assessing global prevalence, 3,011 publicly available metagenomes containing >50 million reads were analyzed (metadata available at https://ftp.ebi.ac.uk/pub/databases/metagenomics/genome_sets/gut_phage_database/GutMetagenomes_metadata.csv). Short reads from each sample were downloaded using SRA Toolkit^103^ v3.0.10, preprocessed using fastp^104^ v0.23.4, and mapped to a database comprising ≥50% complete SPMP and reference vOTUs constructed as described above. Secondary alignments were removed and coverage was estimated using samtools^105^ v1.18. vOTU and VFC detection were performed as described above.

### Host association

Individual viral sequences were linked to host MAGs, and host associations were aggregated for each vOTU. CRISPR arrays were identified from the metagenomic assemblies using CRISPRCasFinder^106^ v4.2.20. Spacers belonging to evidence level 3 and 4 CRISPR arrays were aligned to viral sequences using blastn (options ‘-evalue 1e-6 -ungapped’). Alignments with at most 2 mismatches over the full length of the spacer were used to link viral sequences to MAGs. Viral sequences binned in a MAG and with a flanking host region of ≥10kbp were linked to the MAG.

### Hi-C library preparation and sequencing

Hi-C libraries were generated using the Phase Genomics ProxiMeta Hi-C v4.0 Kit following the manufacturer’s protocol^107^. Faecal material was crosslinked for 15 min at ambient temperature with end-over-end mixing in 1mL of ProxiMeta Crosslinking Solution. The crosslinking reaction was quenched by adding quenching solution and incubating for 20 min at ambient temperature with end-over-end mixing. The sample was washed once with 1× Chromatin Rinse Buffer (CRB), resuspended in 700μL of ProxiMeta Lysis Buffer 1, mixed with 500μL of glass beads, and vortexed for 20 min. Large particles were removed by low-speed centrifugation, and the crosslinked DNA-containing supernatant was transferred to a new tube. A second high-speed centrifugation step was performed to pellet the nuclear fraction, which was washed with 1× CRB. The nuclear pellet was then resuspended in 100μL of ProxiMeta Lysis Buffer 2 and incubated at 65°C for 15 min. Crosslinked DNA was captured on Recovery Beads for 10 min at ambient temperature, followed by magnetic separation and washing with 200μL of 1× CRB.

Crosslinked DNA-bound beads were resuspended in 150μL of ProxiMeta Fragmentation Buffer, and 11μL of ProxiMeta Fragmentation Enzyme was added. Fragmentation was performed for 1 hour at 37°C. After washing the beads with 1× CRB, they were resuspended in 100μL of ProxiMeta Ligation Buffer supplemented with 5μL of Proximity Ligation Enzyme. The proximity ligation reaction was incubated at ambient temperature for 4 hours with end-over-end mixing. 5μL of Reverse Crosslinking Enzyme was added, and the reaction was incubated at 65°C for 1 hour. The DNA was captured on Recovery Beads, Hi-C junctions were bound to streptavidin beads, and the streptavidin-bound Hi-C DNA was washed to remove unbound DNA.

For shotgun library generation, DNA was extracted with the ZymoBIOMICS MagBead DNA kit (D4306) according to the manufacturer’s instructions, and the library was prepared with Watchmaker DNA Library kit according to the manufacturer’s instructions. Library fragment size and concentration were determined using TapeStation and quantitative PCR. Library performance was first determined by low-depth sequencing on an Illumina iSeq System, generating ∼200,000 read pairs. The reads were mapped to the metagenomic assembly and analysed with Phase Genomics’ open-source QC tool, hic_qc. Libraries receiving a ‘Sufficient’ quality judgement from hic_qc proceeded to production sequencing on an Illumina NovaSeq X Plus 10B platform generating paired-end 150bp reads. One library failed QC, leading to the generation of 84 new Hi-C libraries with a minimum and median sequencing depth of 8.1Gbp and 22.6Gbp, respectively (n=108 including 24 libraries previously generated^37^).

### Hi-C–based host association

Hi-C reads were preprocessed using fastp^104^ and mapped to the metagenomic assemblies using bwa-mem (options ‘-5SP’). Reads were filtered to discard unmapped pairs, and secondary and supplementary alignments, using samtools^108^ v1.18 (‘samtools view -F 2316’). Reads were additionally filtered to retain ≥95% identity alignments over ≥80% of the read length.

To distinguish true virus-host interactions from background, a mathematical model for noise in Hi-C data was constructed based on the assumption that Hi-C linkages between MAGs derive primarily from non-specific ligation of extracellular DNA. In this model, different MAGs s form different proportions, *p_s_*, of the extracellular DNA pool, and have different probabilities, *e_s_*, of linking with extracellular DNA. Let *N_s_* be the number of Hi-C read pairs with one end mapping to *s*. Then, the number of reads linking *s* with any other MAG is *M_s_* = *N_s_ e_s_*(1- *p_s_*), and the number of reads linking MAGs *s*_1_ and *s*_2_ is 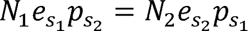. These imply that 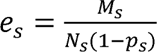 and 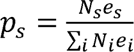. This coupled system of equations for all MAGs s may be solved iteratively using an expectation-maximization–like procedure to obtain parameters for each MAG on a per-sample basis. Following this, p-values for the number of linkages between each viral contig and MAG were computed based on this noise model. In particular, the expected number of Hi-C read pairs linking a viral contig v and a MAG s is 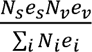. This can be interpreted as a binomial distribution with *N_v_e_v_* ≈ *M_v_* trials and success probability *p_s_*. Then, the p-value is defined as the probability of having at least the observed number of Hi-C linkages between the virus and MAG. P-values within each sample were adjusted for multiple testing using the Benjamini-Hochberg method.

To further eliminate potential false positives, Hi-C linkages between viral sequences and MAGs with adjusted p-value<0.05 were compiled. Because ground truth associations are unknown, VFCs with known host phyla were inspected (*Crassvirales*, *Flandersviridae*, *Konodaiviridae*, *Toyamaviridae*, and *Shinjukuviridae* are predicted to infect Bacteroidota; *Heliusvirales*, *Tsurumiviridae*, and *Okuboviridae* are predicted to infect Firmicutes). Associations with phyla other than the predicted one were taken to be “false”. These linkages might have arisen, for example, from encapsidated phage DNA and extracellular MAG DNA^52^, or by host uptake of viral DNA^109^. A decision tree was constructed using the following variables: number of linkages, adjusted p-value, and the fractions of viral and MAG self- and nonself-linkages. The “false” class was given a class weight of 100. Decision boundaries were established at 2.8% viral self-linkages, followed by 0.065% MAG nonself-linkages. The final filtering criteria were chosen to be: (1) adjusted p-value<0.05, (2) >3% viral self-linkages, (3) >0.1% MAG nonself-linkages, and (4) supported by >1 Hi-C linkage.

### Viral enrichment and virus-like particle (VLP) DNA extraction

Viral enrichment from stool samples and VLP DNA extraction were performed following Shkoporov et al.^110^ with slight modifications. Briefly, 0.5g of fecal material was mixed with 10mL of SM buffer and homogenized by vortexing for 5 min. The suspension was placed on ice for 5 min and clarified by centrifugation at 5,000× g for 10 min at 4°C. The supernatant was transferred to a new tube, subjected to a second centrifugation under the same conditions, and filtered through a 0.45µm vacuum filter (Sartorius, 180F01-E8). Sodium chloride (NaCl; Sigma-Aldrich, S9888-1KG) and poly(ethylene glycol) (PEG-8000; Sigma-Aldrich, 89510-1KG-F; final concentrations 0.5M and 10% w/v, respectively) were added to the filtrate and mixed until fully dissolved. Samples were then incubated overnight at 4°C to allow VLPs to precipitate.

Precipitated VLP pellets were collected by centrifugation at 10,000× g for 20 min at 4°C and supernatants were discarded. Excess liquid was removed by inverting the tubes on a paper towel for 5 min. Pellets were resuspended with 400µL of SM buffer and extracted with equal volumes of chloroform (Kanto, 07278-00). Phase separation was achieved by centrifugation at 2,500× g for 5 min and the aqueous layer was transferred to microcentrifuge tubes. A 40µL solution of 10mM calcium chloride (CaCl_2_; Sigma-Aldrich, C1016-100G) and 50mM magnesium chloride (MgCl_2_; Sigma-Aldrich, 208337-100G) was added, followed by 8U TURBO DNase (Invitrogen, AM2239) and 20U RNase A (QIAGEN, 19101). Samples were incubated at 37°C for 1 hour, then heated to 70°C for 10 min to inactivate the enzymes.

40µg of Proteinase K (QIAGEN, 19134) and 20µL of 10% sodium dodecyl sulfate (SDS) (Invitrogen, AM9820) were then added, and the samples were incubated at 56°C for 20 min. Buffer RLT (QIAGEN, 79216) was added and samples were incubated at 65°C for 10 min to lyse the VLPs. The lysate was extracted twice using equal volumes of Phenol:Chloroform:Isoamyl Alcohol (25:24:1; Sigma-Aldrich, P2069-400ML), centrifuging each time at 8,000× g for 5 min at room temperature. The final aqueous phase was purified using Qiagen DNeasy Blood & Tissue Kit (QIAGEN, 69506) according to the manufacturer’s instructions and eluted in 50µL. DNA was quantified using a Qubit 4 Fluorometer (Thermo Scientific^TM^, Q33238).

### VLP DNA library preparation and sequencing

VLP DNA library preparation was performed with the NEBNext^®^ Ultra^TM^ II FS DNA Library Prep Kit (New England Biolabs, E7805) according to the manufacturer’s instructions. Barcoding and enrichment were done using the NEBNext^®^ Multiplex Oligos for Illumina^®^ (96 Unique Dual Index Primers Pairs; New England Biolabs, E6440). Paired-end sequencing (2×150bp reads) was performed on the Illumina NovaSeq X. A total of 64 VLP sequencing libraries were generated, with one sample having a depth of 71Mbp and the others a minimum and median depth of 1.9Gbp and 5.8Gbp, respectively. Illumina reads were preprocessed using fastp^104^ v0.23.4. The degree of viral enrichment of each sample was estimated as the ratio of read recruitment proportion to the SILVA^111^ release 138.1 SSU Ref NR99 database (at ≥95% identity over ≥80% of the read length) for the bulk versus VLP sample.

### Detection of actively replicating phages

Four separate metrics were used to identify replicating vOTUs in a sample. The first two are the vOTU/host MAG mean coverage ratio in the bulk and VLP sample, respectively. Short reads from each sample were mapped to the MAGs assembled from that sample, and reads were filtered to retain properly paired mappings at ≥95% identity over ≥80% of the read length. The mean coverage of each MAG was computed excluding the top and bottom 10% of base coverage values. A vOTU/MAG coverage ratio >1.75 was used as a replication signature^59^.

The other two metrics rely on the mean coverage of *non-viral* contigs as a null distribution. Short reads from each sample were mapped to all contigs assembled from that sample, and reads were filtered to retain properly paired mappings at ≥95% identity over ≥80% of the read length. Only contigs ≥5kbp that do not contain any viral regions were retained. A non-viral contig was considered present in a sample if reads cover ≥70% of its length, following the vOTU detection threshold. The mean coverage of each non-viral contig was computed excluding the top and bottom 10% of base coverage values. vOTUs whose mean coverages were outliers compared to the null distribution, i.e. exceeded the median + 2×inter-quartile range of non-viral contigs in the VLP sample, were flagged as having a replication signature.

Finally, to combine information from the bulk and VLP samples in a consistent way, the reads per kilobase million (RPKM) of each vOTU and non-viral contig was calculated, given by the number of reads mapped (with a pseudocount of 1 added) divided by the sequence length (in kbp) and the number of million reads in the sequencing library. vOTUs whose VLP/bulk RPKM ratio were outliers, i.e. exceeded the median + 2×inter-quartile range of non-viral contigs, were flagged as having a replication signature.

### Bacterial strains and culture conditions

Genome assemblies of publicly available bacterial strains were downloaded, and prophages were identified using geNomad^41^. Prophages were aligned with vOTUs using blastn (options ‘-evalue 1e-12 -perc_identity 95 -max_target_seqs 10000’). Alignments between pairs of sequences were merged using ‘anicalc.py’ from the CheckV package. Alignments covering ≥60% of both prophage and vOTU were checked. By this criterion, *Ruminococcus torques* Holdeman and Moore (ATCC 22756), *Ruminococcus gnavus* Moore et al. (ATCC 29149), *Anaerostipes hadrus* (ATCC 29173), and *Agathobacter rectalis* (Hauduroy et al.) Prevot (ATCC 33656) carry VFC 1, 2, 4, and 4 prophages, respectively. These strains were procured for follow-up investigations.

Bacterial strains were propagated using Brain Heart Infusion (BHI) broth (Thermo Scientific^TM^, CM 1135B) supplemented with 0.05% L-cysteine hydrochloride monohydrate (Sigma-Aldrich, C6852-100G) and 0.02% dithiothreitol (DTT) (Thermo Scientific^TM^, R0862). All culture media were pre-reduced in an anaerobic chamber (atmosphere of N_2_ (75%), CO_2_ (20%), and H_2_ (5%)) for at least 24 hours. For bacterial DNA extraction, 1mL of overnight culture was centrifuged at 8,000× g for 5 min, after which the supernatant was removed. The resulting bacterial pellet was resuspended in 800µL of C1 buffer from the Qiagen DNeasy PowerSoil Pro Kit (Qiagen, 47016). Samples were then transferred to PowerBead Pro Tubes, and DNA extraction was performed according to the manufacturer’s instructions, with a final elution volume of 100µL. Full-length 16S rRNA was amplified by PCR using primers from Matsuo et al.^112^ (**Supplementary Table 2**). The PCR amplicons were purified using 1.5× AMPure XP beads (Beckman Coulter, A63882) prior to Sanger sequencing at Axil Scientific. Sequences were then BLAST-ed to confirm strain identity.

### Prophage induction and purification of VLPs

Overnight bacterial cultures were inoculated into 40mL of BHI media to an OD_600nm_ of 0.05, mixed, and grown in the anaerobic chamber statically at 37°C until the cultures reached mid-log phase (OD_600nm_=0.4–1.25). Different amounts of mitomycin C (mitC; Fisher Scientific, FSKJ67460.XF; 0, 0.3, 1.0, and 5 µg/mL) were added to the culture, inverted several times, and incubated in the anaerobic chamber overnight at 37°C. To obtain higher phage titer, prophage induction and purification of VLPs were done in larger volumes using the previously described protocol with slight modifications. Briefly, 20mL of overnight bacterial culture was added to 2L of media and incubated over 3 days at 37°C in the anaerobic chamber.

Cultures were centrifuged at 5,000× g for 10 min, before filtering the supernatant through a 0.45µm vacuum filter (Sartorius, 180F01-E8). NaCl and PEG-8000 (final concentrations 0.5M and 10% w/v, respectively) were added to the filtrate and dissolved completely, and the samples were incubated overnight at 4°C. The precipitate was then centrifuged at 10,000× g for 1 hour at 4°C, and the supernatant was gently removed. The pellet was air-dried for 5 min and resuspended in 400µL of SM buffer, followed by the addition of 40µL chloroform. The sample was mixed by inversion and centrifuged at 2,500× g for 5 min. The aqueous phase was transferred to a new tube and filtered through a 0.45µm filter. The filtrate was kept at 4°C for short-term storage and at −80°C for long-term storage.

### Validation of prophage induction

VLP samples were treated with TURBO DNase™ (Invitrogen™, AM2238) following the manufacturer’s instructions with slight modifications. Briefly, 50µL of the VLP sample was aliquoted and 5 µL of 10× Turbo DNase buffer, 1µL of Turbo DNase, and 1µL of RNase A were added. Samples were incubated at 37°C for 1 hour before enzyme inactivation at 70°C for 10 minutes.

PCR primers were designed using SnapGene v8.0.1 to target the phage region (PR) and the junction of circularization of the phage (JoC; **Supplementary Table 3**). Bacterial genomic DNA (gDNA) and DNase-treated VLP samples were used as templates, with water as negative control. Amplification was performed using Phusion High Fidelity PCR Master Mix with HF Buffer (Thermo Scientific™, F531L) with the following conditions: initial denaturation at 98°C for 1 min, 35 cycles of 98°C for 15 sec, primer-specific annealing temperature for 15 sec and 72°C for 30 sec, followed by a final extension at 72°C for 10 min. Up to 50ng of PCR products were visualized on a 1.5% agarose gel run at 120V for 40 min in TAE buffer, alongside a 1kbp DNA ladder (Promega, G5711).

For sequencing of induced phages, viral enrichment and VLP DNA extraction were performed as previously described. MitC-treated samples were pooled together, and library preparation of the samples with and without mitC treatment was performed following the previously described protocol and sequenced on an Illumina NovaSeq X Plus system. Illumina reads were preprocessed using fastp^104^, mapped to the isolate genome assemblies using bwa-mem, and processed further using samtools (‘samtools depth -H’). Packaging strategies of the phages were inferred using PhageTerm^113^ v1.0.12.

### Phage qPCR and transmission electron microscopy imaging

To quantify phage titer and ensure sufficient concentration for transmission electron microscopy (TEM) imaging, qPCR was performed on the DNase-treated VLP samples using PowerUp™ SYBR™ Green Master Mix (Applied Biosystems, A25742; primers and g-blocks listed in **Supplementary Table 4**). VLP samples were fixed using EM-grade glutaraldehyde solution (Sigma-Aldrich, G5882) to a final concentration of 1% at room temperature for 20 minutes. Carbon film-supported copper grids (EMCN, 300 mesh) were glow-discharged for 60 sec, after which 5µl of the fixed sample was applied onto the grid and incubated at room temperature for 5 minutes. The sample was then blotted away and washed once with Milli-Q water to wash off excess buffer. The grid was stained with 5µl of UranyLess EM Stain (Electron Microscopy Sciences, 22409) for 30 sec. The stain was immediately blotted away, and the staining step was repeated once, after which the grid was blotted completely and air-dried. Electron micrographs were acquired using an FEI Tecnai T12 transmission electron microscope (Thermo Fisher Scientific) equipped with a LaB6 filament, operated at an accelerating voltage of 120kV, and recorded with a 4k CCD camera.

### Functional annotation of viral genomes

Diversity generating retroelements (DGRs) were identified in the ≥50% complete vOTUs using DGRscan^114^. Antiphage defence and anti-defense systems were identified in the ≥50% complete vOTUs using DefenseFinder^115^ v2.0.0 (options ‘--antidefensefinder’). Auxiliary metabolic genes (AMGs) were identified in the vOTUs by running VirSorter2^39^ (options ‘--prep-for-dramv’) followed by DRAM-v^116^ v1.5.0. Circos plots were drawn using the pyCirclize v1.10.1 package.

### Single nucleotide variant (SNV) and pN/pS analysis

To identify SNVs in vOTUs, a conservative mapping approach was developed. A MAG database was created by dereplicating all SPMP MAGs at 99% identity using dRep^117^ v3.4.2 (options ‘-pa 0.99 --ignoreGenomeQuality’) and removing prophage regions using seqtk v1.4. Illumina reads were mapped to a combined database of vOTUs and dereplicated MAGs, and filtered to retain properly paired mappings at ≥95% identity over ≥80% of the read length. vOTUs with ≥70% coverage breadth were kept for further analysis. SNVs were called using LoFreq^118^ v2.1.5, and SNVs with AF=1 were excluded to retain only positions with multiple variants in a sample.

SNVs were classified as genic or intergenic, and genic SNVs classified as synonymous, nonsynonymous, or in a stop codon for each gene they appear in. The number of possible synonymous and nonsynonymous SNVs was computed by substituting each codon position by each of the 3 alternative nucleotides (3×3=27 possible variants per codon) and summing across all codons in a gene (codons containing non-standard characters and stop codons were excluded). A pseudocount of 1 was added to the observed counts for each gene. The pN/pS ratio of a gene was computed as the ratio of observed over possible nonsynonymous SNVs, divided by the ratio of observed over possible synonymous SNVs.

### Prevalence modelling of VFCs

All VFCs with size ≥10 were used for modelling prevalence in the SPMP cohort. Eleven predictors were considered: median VFC abundance, host phylum, family and order entropies, median host abundance, and the percentages of vOTUs that are temperate, encode DGRs, carry AMGs, carry defense systems, carry anti-defense systems, and replicate in at least one sample. For generalized linear modelling, all numerical predictors were first normalized using min-max scaling. The fractional VFC prevalence was modelled with Elastic Net regularization (α=0.5) and 10-fold cross-validation using the glmnet package^119^ v4.1.10. For random forest regression, categorical variables were one-hot encoded, and hyperparameter optimization performed using 5-fold cross-validation and R^2^ as the scoring metric. Feature importances were averaged across folds to identify the most influential predictors.

## Data availability

All Supplementary Data files (**Supplementary Data 1–11**), identified viral sequences, and representative vOTU sequences are available via Zenodo at DOI:10.5281/zenodo.18253940. Hi-C and VLP metagenomic sequencing reads are available from the European Nucleotide Archive (ENA) under project accession PRJEB106095.

## Code availability

All code used to generate the results and figures in this study are available at https://github.com/CSB5/SPMP_Phages.

## Acknowledgements

The authors thank E. Adriaenssens and R. Krishnamurthy for critical reading of a draft of this manuscript. This work was supported by a National Medical Research Council Open Fund – Large Collaborative Grant (grant no. 23-0614). NN was supported by a National Research Foundation Investigatorship grant (grant no. NRFI09-0015). HC was supported by an A*STAR Career Development Fund (project no. C210812044). This work was supported by the A*STAR Computational Resource Centre through the use of its high performance computing facilities.

