## supplementary figures and tables for "GuFi phages represent the most prevalent viral family-level clusters in the human gut microbiome"

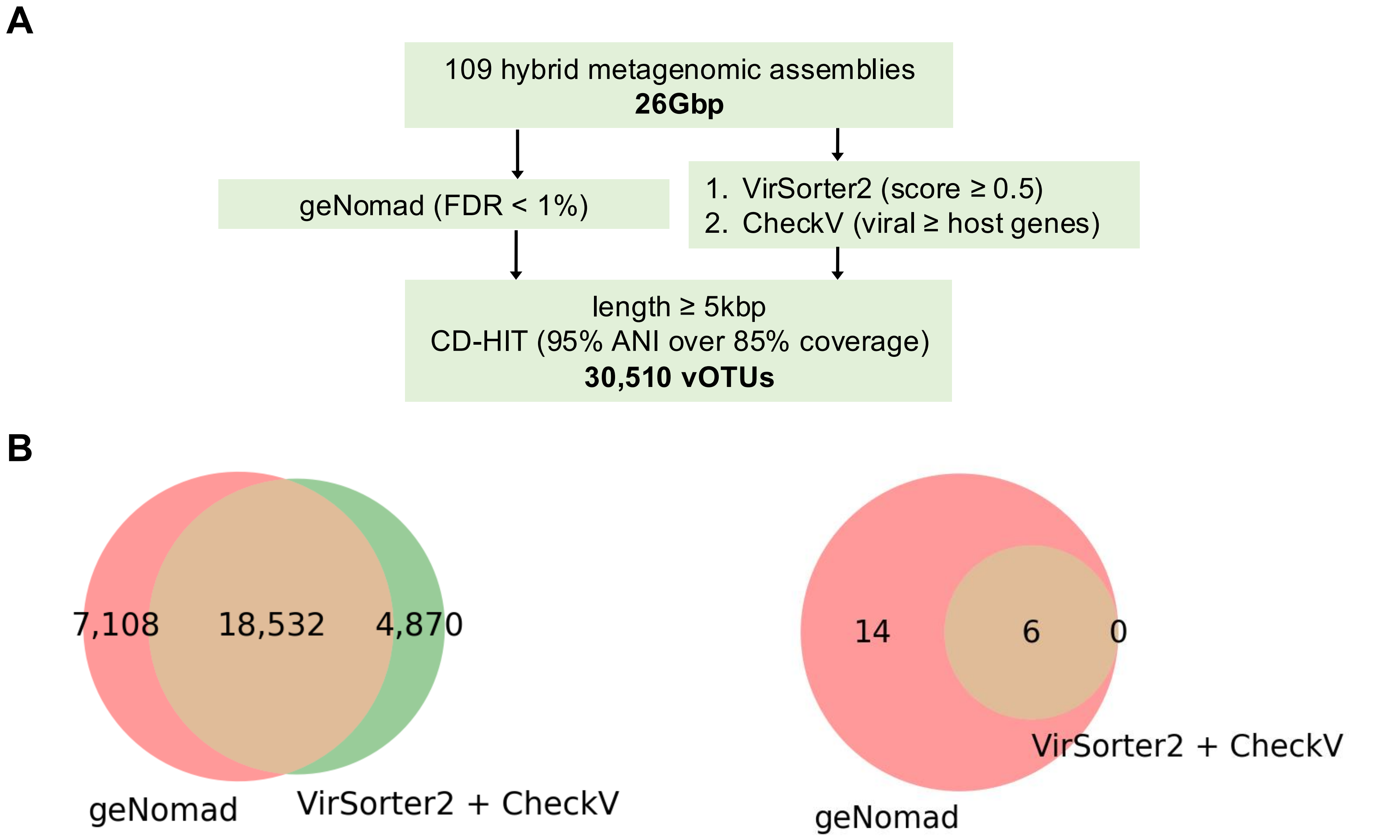


**<u>Supplementary Figure 1:</u> Identification of vOTUs from hybrid metagenomic assemblies.** (A) Pipeline used to identify viral sequences and cluster them into vOTUs. (B) geNomad and VirSorter2 + CheckV approaches identify complementary sets of vOTUs. Number of vOTUs (left) and jumbo (≥200kbp) vOTUs (right) recovered using either or both approaches.

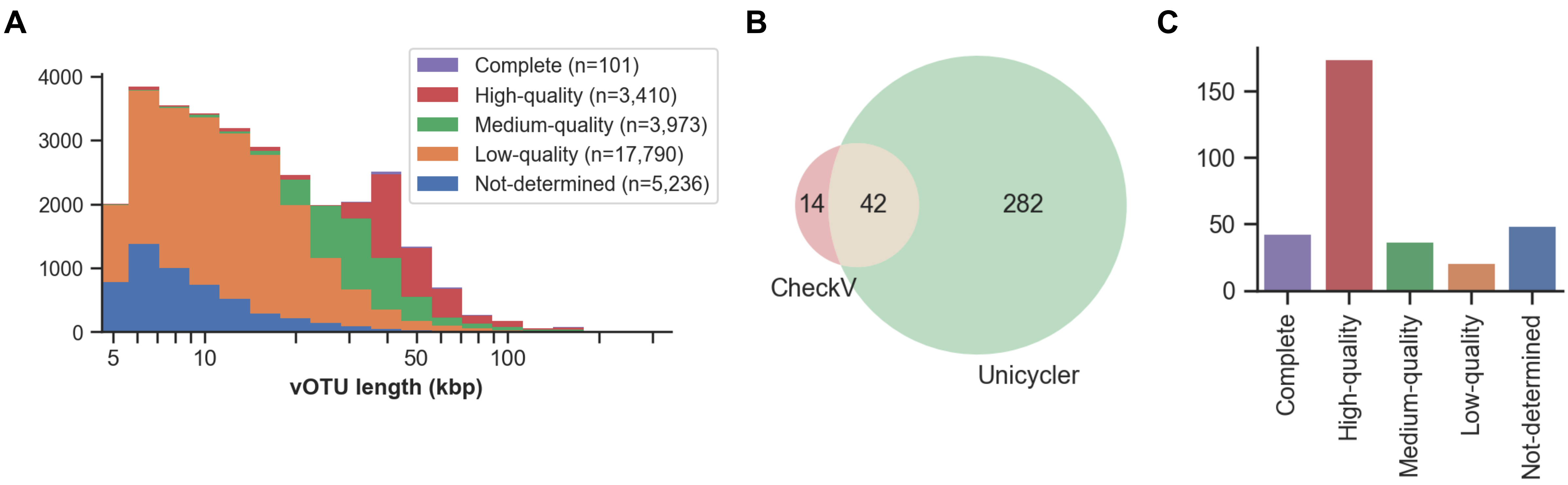


**Supplementary Figure 2: Recovery of 3,511 high-quality and complete vOTUs.** (A) vOTU length distribution color-coded by CheckV quality tiers. A total of 3,973 were estimated to be 50-90% complete ('medium-quality'), 3,410 as ≥90% complete ('high-quality'), and 101 as 'complete'. (B) Number of circular vOTUs as assessed by CheckV, having a circular Unicycler assembly, or both. Most of the circular assemblies were missed by CheckV, potentially due to terminal repeats not being present in the assembly. (C) CheckV quality tier distribution of vOTUs having a circular Unicycler assembly (n=324), with the majority estimated as 'high-quality'. CheckV may also overlook 'complete' viral genomes whose flanking host regions were removed by the viral contig identification tool.

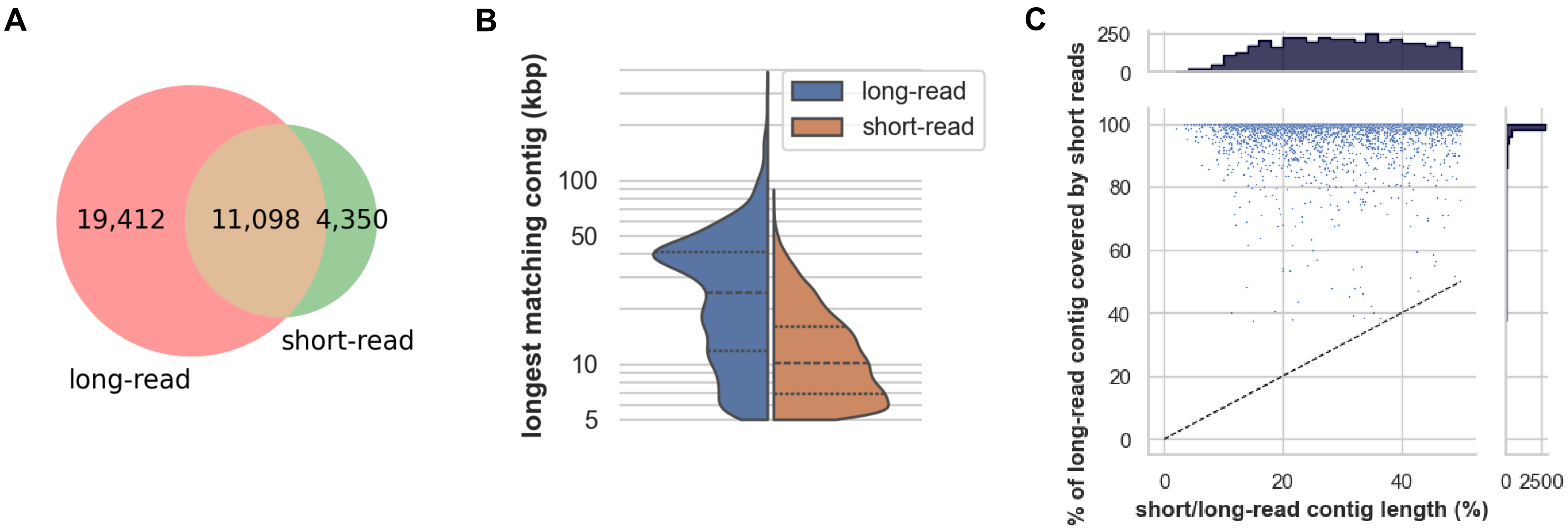


**<u>Supplementary Figure 3:</u> Viral recovery is significantly improved using long-read versus short-read assembly.** (A) Long-read and short-read contigs were jointly dereplicated (**Methods**), and the number of clusters containing long-read contigs only, short-read contigs only, or both are shown. (B) Distribution of the longest long-read and short-read contig in clusters containing both (n=11,098, median of 24.7kbp for long-read and 10.2kbp for short-read). (C) Short reads were mapped to long-read contigs that are >2× longer than the longest short-read contig in the same cluster (n=3,911). Mapped reads were filtered to have ≥99% identity over ≥80% of the read length. The vast majority of long-read contigs have high short-read coverage (median of 99.6%), suggesting that incomplete short-read viral recovery was due to more fragmented assemblies rather than regions that are only sequenced by long reads.

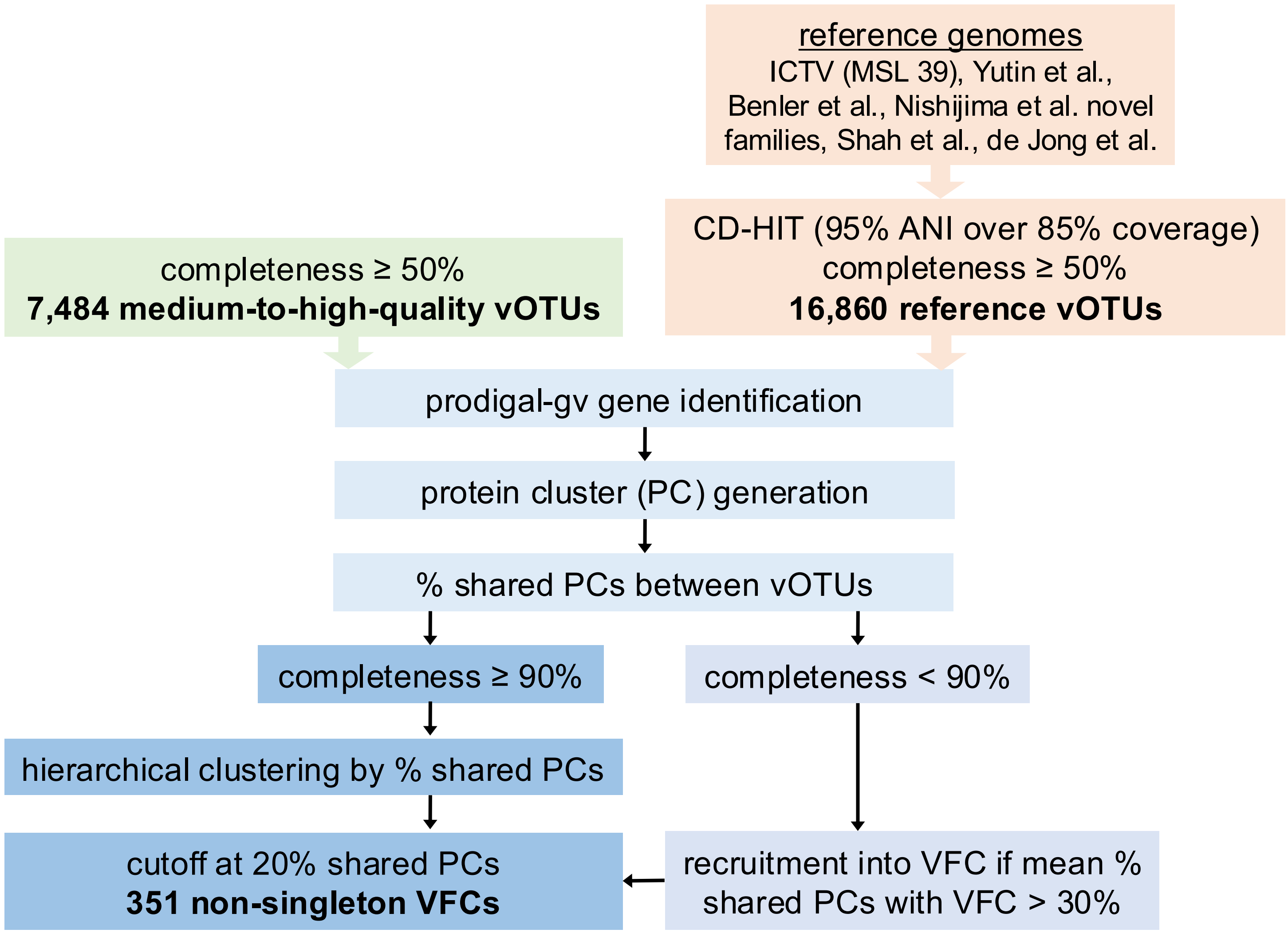


**Supplementary Figure 4: Construction of VFCs from SPMP vOTUs and reference genomes.** Viral family-like clusters (VFCs) were constructed from medium and high-quality (≥50% complete) and complete vOTUs and reference genomes. Protein sequences were combined and protein clusters (PCs) were generated. Pairwise percentage shared PCs were computed, and high-quality (≥90% complete) and complete vOTUs were hierarchically clustered by percentage shared PCs. Clustering of reference families were assessed at different cutoffs, leading to the formation of VFCs at a cutoff of 20% shared PCs. Medium-quality (50-90% complete) vOTUs were recruited if the mean percentage shared PCs with high-quality vOTUs exceeds 30% (**Methods**). This resulted in the creation of 351 non-singleton VFCs (containing at least two SPMP vOTUs).

| VFC | 1 | 13 | 16 | 43 | 49 | 242 | 117 | 67 | 123 | 142 | 47 | 90 | 63 | 65 | 546 | 108 | 109 | 218 | 125 | 326 | 183 | 220 | 243 |
|---|---|---|---|---|---|---|---|---|---|---|---|---|---|---|---|---|---|---|---|---|---|---|---|
| *Heliusvirales* | 712 (95) | | | | | | | | | | | | | | | | | | | | | | |
| *Tsurumiviridae* | | 85 (99) | | | | | | | | | | | | | | | | | | | | | |
| *Okuboviridae* | | | 70 (100) | | | | | | | | | | | | | | | | | | | | |
| *Konodaiviridae* | | | | 64 (100) | | | | | | | | | | | | | | | | | | | |
| *crAssphage-α* | | | | | 132 (100) | | | | | | | | | | | | | | | | | | |
| *crAssphage-β* | | | | | | 71 (99) | | | | | | | | | | | | | | | | | |
| *crAssphage-γ* | | | | | | | 60 (100) | | | | | | | | | | | | | | | | |
| *crAssphage-δ* | | | | | | | | 217 (99.5) | | | | | | | | | | | | | | | |
| *crAssphage-ε* | | | | | | | | | 30 (100) | | | | | | | | | | | | | | |
| *crAssphage-ζ* | | | | | | | | | | 69 (100) | | | | | | | | | | | | | |
| *Peduoviridae* | | | | | | | | | | | 103 (83) | 21 (17) | | | | | | | | | | | |
| *Toyamaviridae* | | | | | | | | | | | | | 75 (97) | | | | | | | | | | |
| *Microviridae* | | | | | | | | | | | | | | 6 (30) | 14 (70) | | | | | | | | |
| *Shinjukuviridae* | | | | | | | | | | | | | | | | 52 (100) | | | | | | | |
| *Aliceevansviridae* | | | | | | | | | | | | | | | | | 113 (91) | 11 (9) | | | | | |
| *Autographiviridae* | | | | | | | | | | | | | | | | | | | 214 (54) | 38 (10) | | | |
| *Flandersviridae* | | | | | | | | | | | | | | | | | | | | | 298 (100) | | |
| *Drexlerviridae* | | | | | | | | | | | | | | | | | | | | | | 254 (100) | |
| *Umezonoviridae* | | | | | | | | | | | | | | | | | | | | | | | 112 (100) |

**Supplementary Figure 5: Agreement of VFCs with known viral families.** Number of reference vOTUs with family-level taxonomic assignments (row) clustered in each VFC (column), with percentages shown in parentheses. All best hits of ICTV-ratified families (underlined) with non-singleton VFCs are shown. VFC clustering of other known families, such as those described in de Jonge et al., Nishijima et al., and Yutin et al., are also shown. Zero entries are left blank. VFCs with >95% of references correctly grouped are highlighted in green. In some cases, VFC reconstruction split known families into two (highlighted in blue). Further inspection revealed, for example, that one of the *Peduoviridae* VFCs only infects Proteobacteria while the other infects Desulfobacterota hosts. The 2 *Microviridae* VFCs correspond to the 2 ICTV-ratified subfamilies, *Bullavirinae* and *Gokushovirinae*. Another example is *Autographiviridae*, which has since been promoted to the order *Autographivirales*, reflecting the still-evolving nature of viral taxonomy.

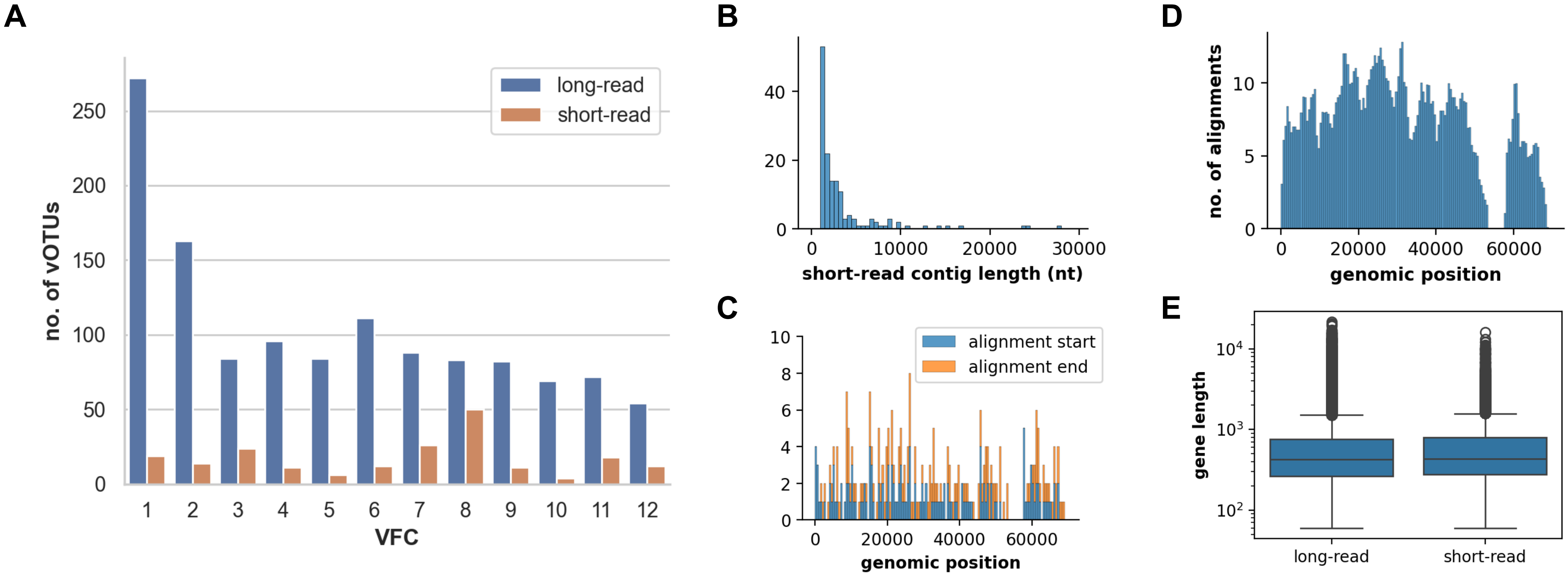


**<u>Supplementary Figure 6:</u> VFC construction from short-read viral contigs fails to reconstruct highly prevalent VFCs.** (A) An identical pipeline was used to construct VFCs from short-read vOTUs and the same set of reference genomes (**Supplementary Figure 4**). Number of vOTUs in the best-hit short-read VFC (orange) are shown alongside the corresponding long-read VFC (blue). Best hits were found by jointly dereplicating short-read and long-read viral contigs (**Supplementary Figure 3**). (B) Histogram of short-read contig lengths aligning to a 'complete' 68kbp VFC 1 vOTU assembled with long reads (based on 99.9% ANI over 99.9% coverage of short-read contig using blastn; '-task megablast -evalue 1e-12', n=147). (C) Stacked histogram of start and end coordinates of short-read contig alignments over the long-read vOTU, binned in windows of 500bp. (D) Mean number of alignments over the long-read vOTU, binned in windows of 500bp. A region between 53.4-57.8kbp along the long-read contig is missing, potentially unassembled in all samples. (E) Gene length distributions for ≥50% complete long and short-read vOTUs, used for protein cluster (PC) generation. The medians are 423 and 432 bp, respectively, a 2% difference.

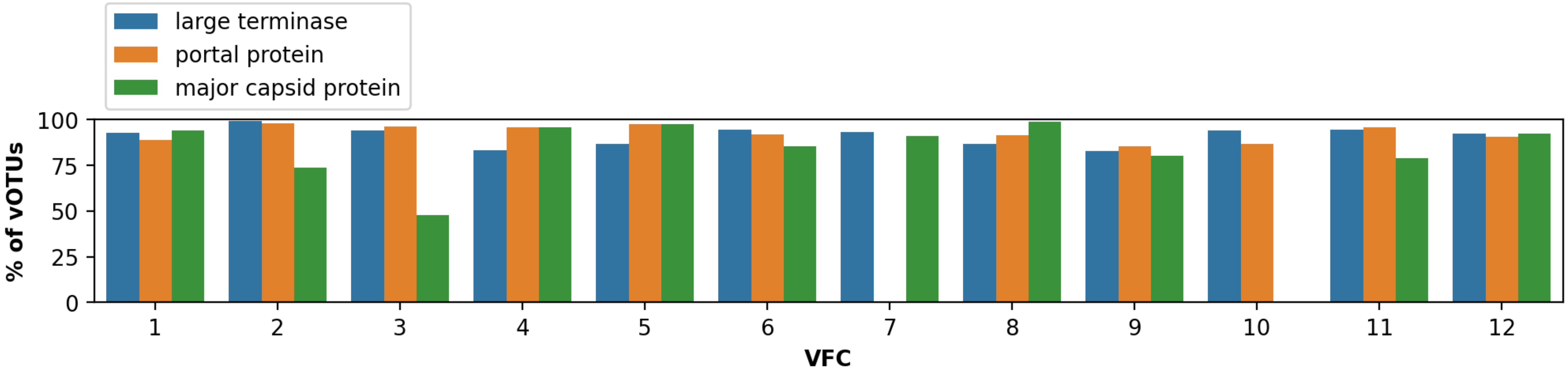


<u>Supplementary Figure 7:</u> **Functional annotation of protein clusters in the top VFCs.** Percentage of vOTUs in the most prevalent VFCs with annotated large terminases (blue), portal proteins (orange), and major capsid proteins (MCP; green) after manual inspection of geNomad and pharokka annotations to assign protein clusters (PCs) to these functions. The lack of matches to structural proteins in some VFCs, for example the portal protein of VFC 7 and the MCP of VFC 10, may be due to protein-level divergence in these clades. Indeed, these VFCs have numerous PCs that are carried by >90% of members but unannotated by either geNomad or pharokka (**Supplementary Data 11**).

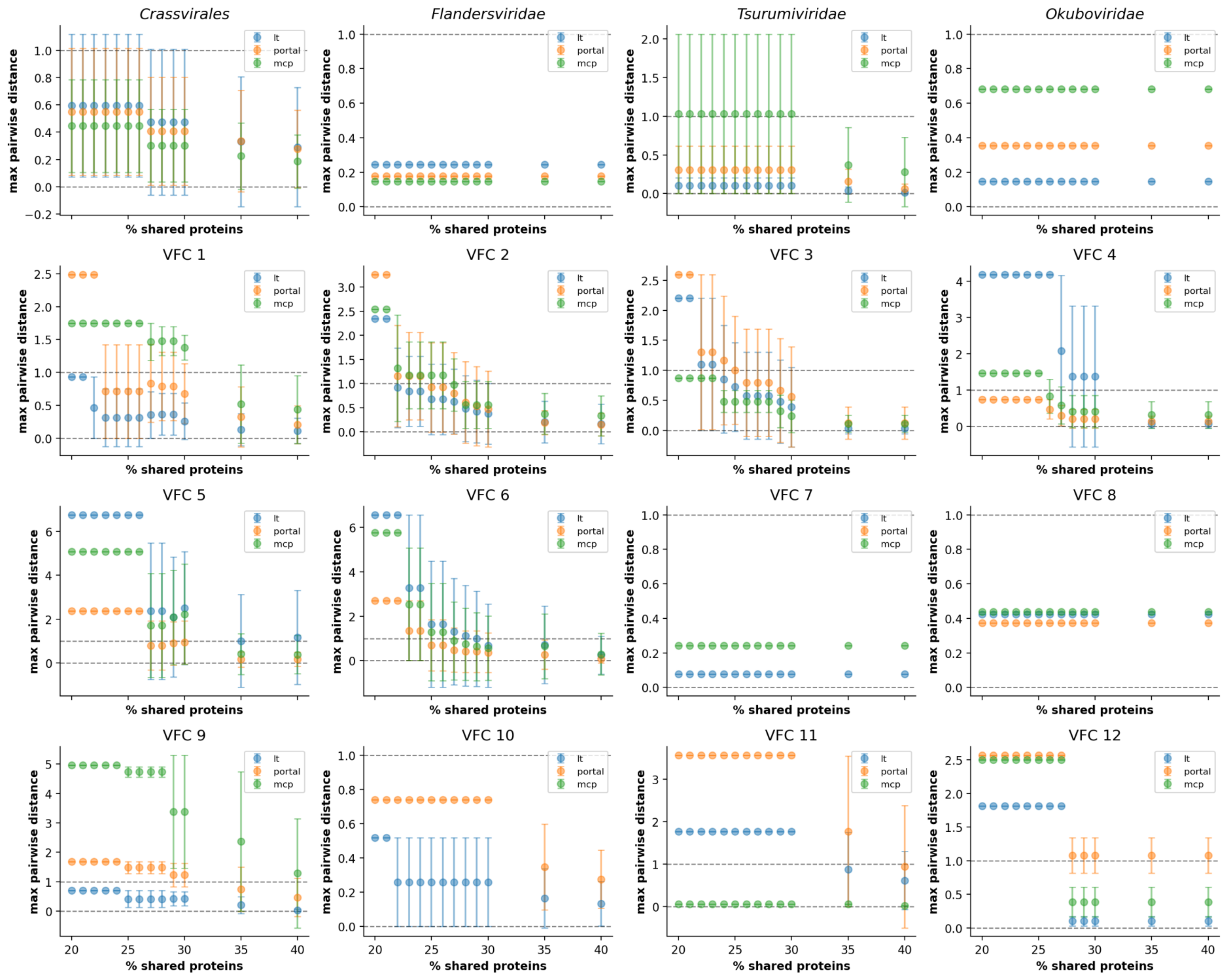


**<u>Supplementary Figure 8:</u> Plateaus in phylogenetic dispersion as a signature that VFCs are robustly defined.** Phylogenetic dispersion (PD), defined as the maximum pairwise phylogenetic distance, of large terminases (blue; lt), portal proteins (orange), and major capsid proteins (green, mcp) in subclades formed by hierarchical clustering at different percentage shared protein cluster (PC) cutoffs, within reference *Crassvirales*, *Flandersviridae*, *Tsurumiviridae*, and *Okuboviridae* genomes (top row) and VFCs 1-12. Up to 3 genes were sampled from each subclade sharing 40% PCs ('genus-level'; **Methods**). Error bars denote the standard deviation in PD among subclades formed at each value of the cutoff.

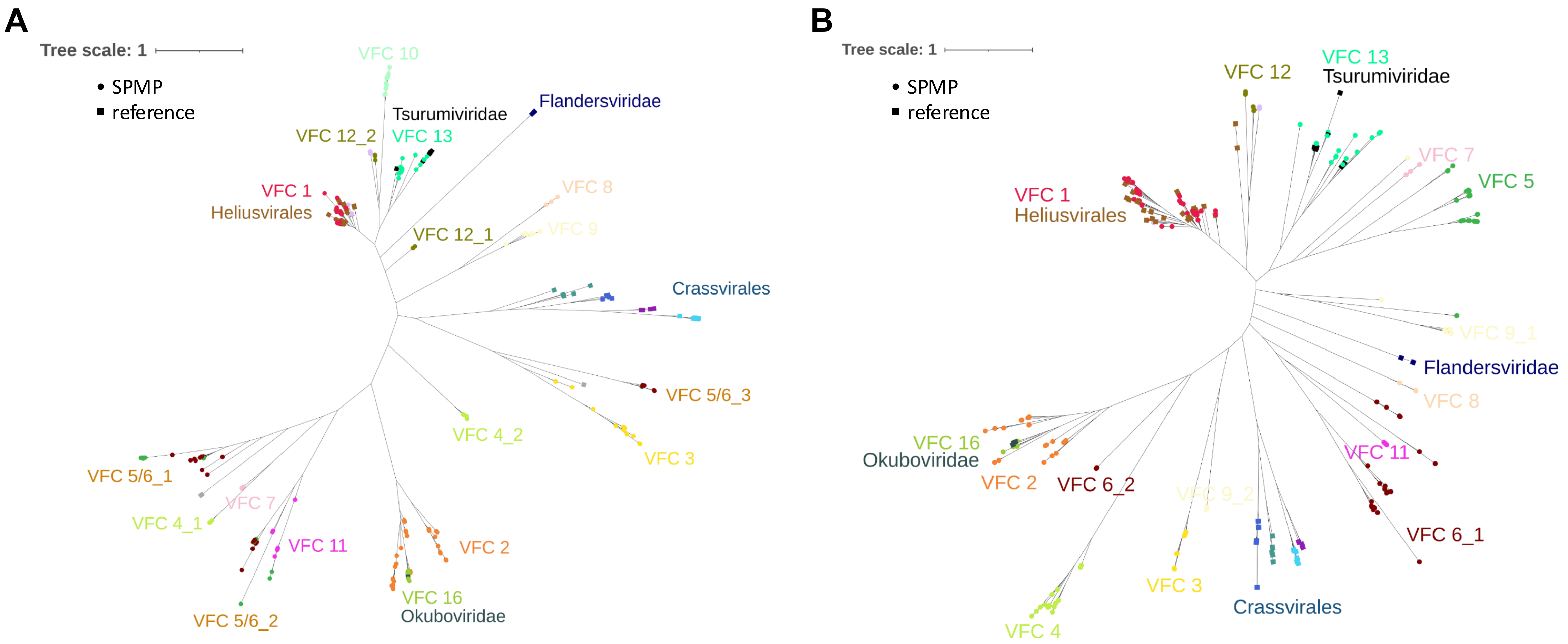


<u>**Supplementary Figure 9:**</u> **Phylogenetic trees of other key viral structural proteins.** Phylogenetic trees for large terminases (A) and major capsid proteins (B) sampled from vOTUs in the prevalent VFCs and reference *Heliusvirales*, *Crassvirales*, *Flandersviridae*, *Tsurumiviridae*, and *Okuboviridae* genomes (**Methods**, **Supplementary Data 4**). Clusters resemble VFC clustering and the portal protein tree (**Fig. 1E**), with some evidence for subfamily-level divergence and potential horizontal gene transfer events. For example, VFC 4 has one portal protein and one MCP cluster but 2 distinct large terminase clusters. Also, VFCs 5 and 6 each have one portal protein cluster but 3 distinct large terminase clusters mixed together.

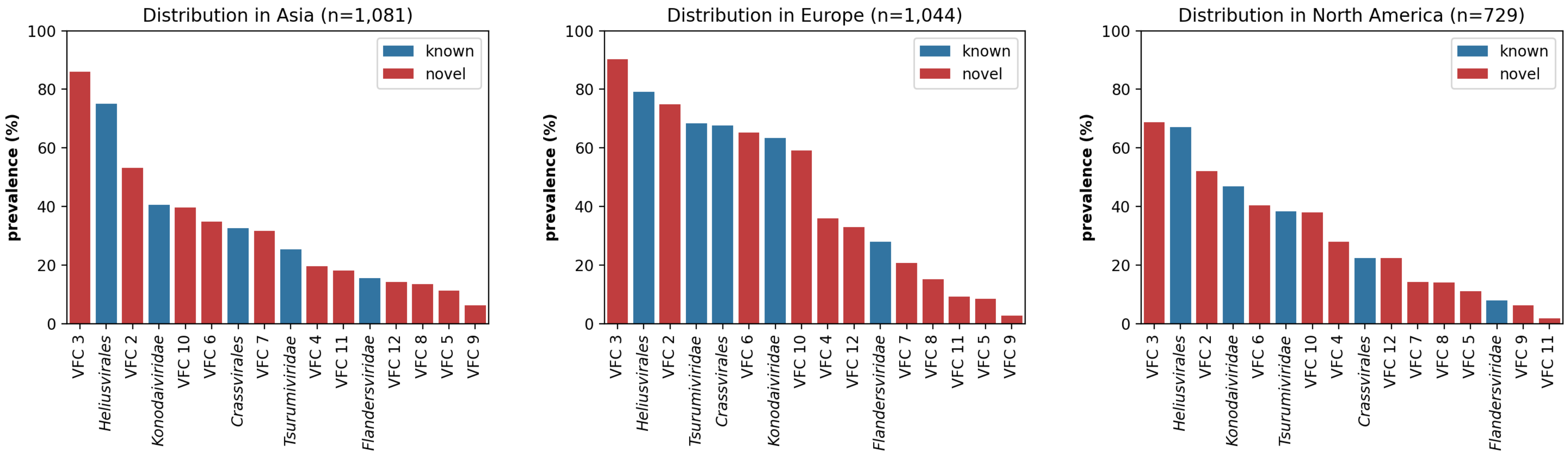


**<u>Supplementary Figure 10:</u> Prevalence of VFCs and reference clades across multiple continents.** Prevalence of top VFCs and selected reference clades in Asia (left), Europe (middle), and North America (right). Samples with >50 million reads were mapped to SPMP and reference vOTUs belonging to these viral groups (**Methods**). A clade is identified as present in a sample if at least 70% of one member vOTU is covered. Several VFCs that are highly prevalent in the SPMP cohort are also highly prevalent in multiple continents, in particular VFCs 3, 1 (*Heliusvirales*), and 2.

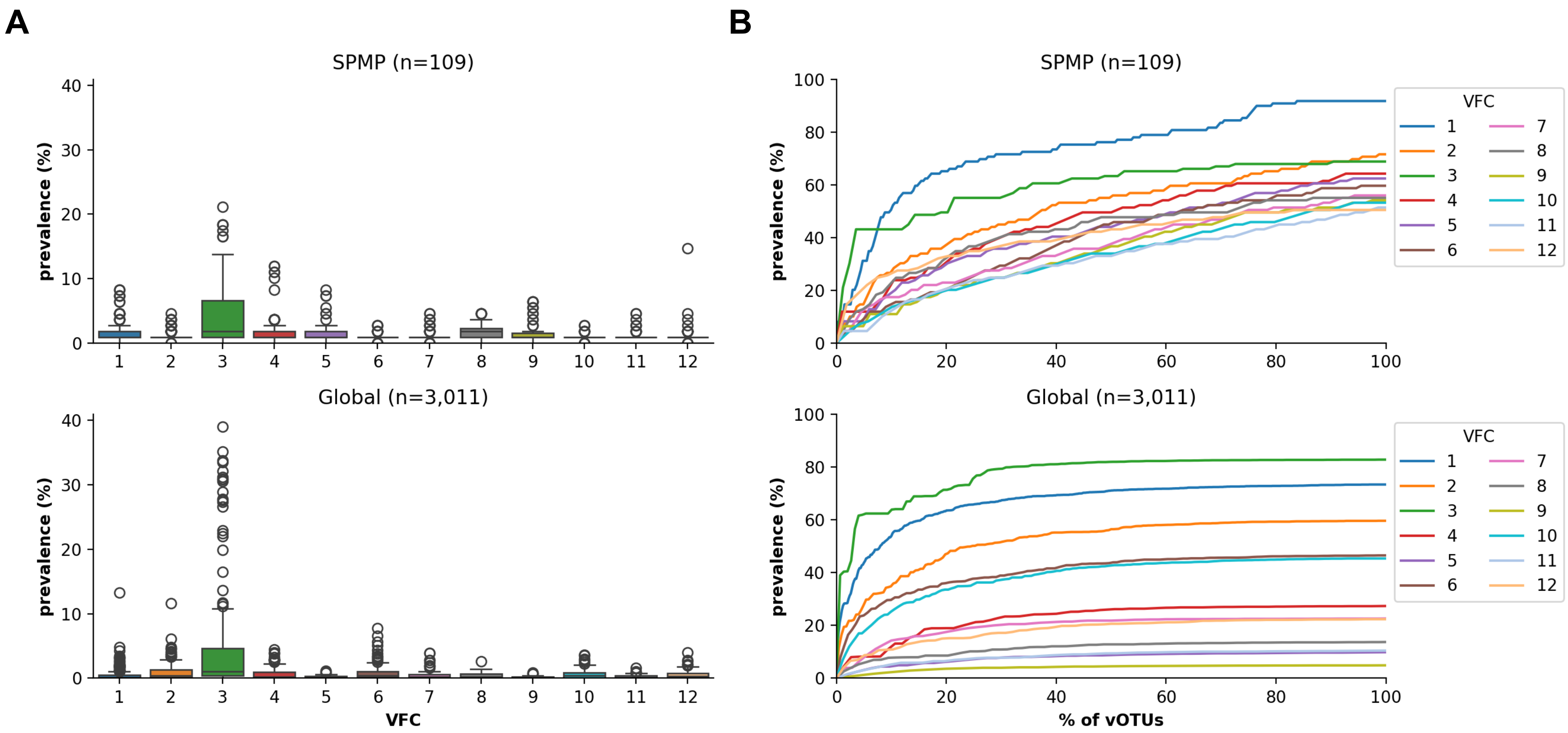


**<u>Supplementary Figure 11:</u> Individual vOTU prevalence does not contribute significantly to overall VFC prevalence, except for VFC 3.** (A) Distribution of vOTU prevalence in the SPMP cohort (top) and globally (bottom). (B) VFC prevalence as vOTUs are added in descending order of vOTU prevalence, in the SPMP cohort (top) and globally (bottom).

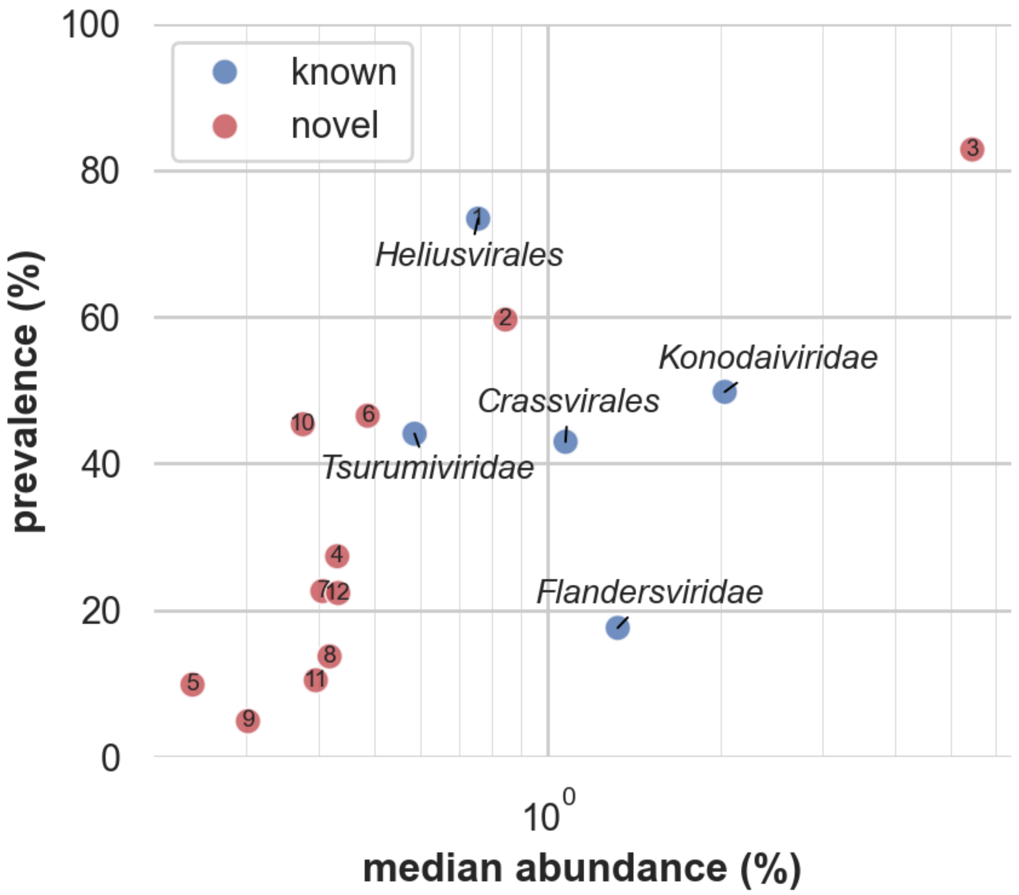


<u>**Supplementary Figure 12:**</u> **Prevalence versus median abundance of VFCs and reference families in global datasets.** Well-known phage families including *Crassvirales* and *Flandersviridae* have high median abundance globally, though VFC 3 appears to be even more abundant and prevalent. Several abundant VFCs in our cohort are less abundant globally (e.g. VFC 8), highlighting the importance of population-specific studies.

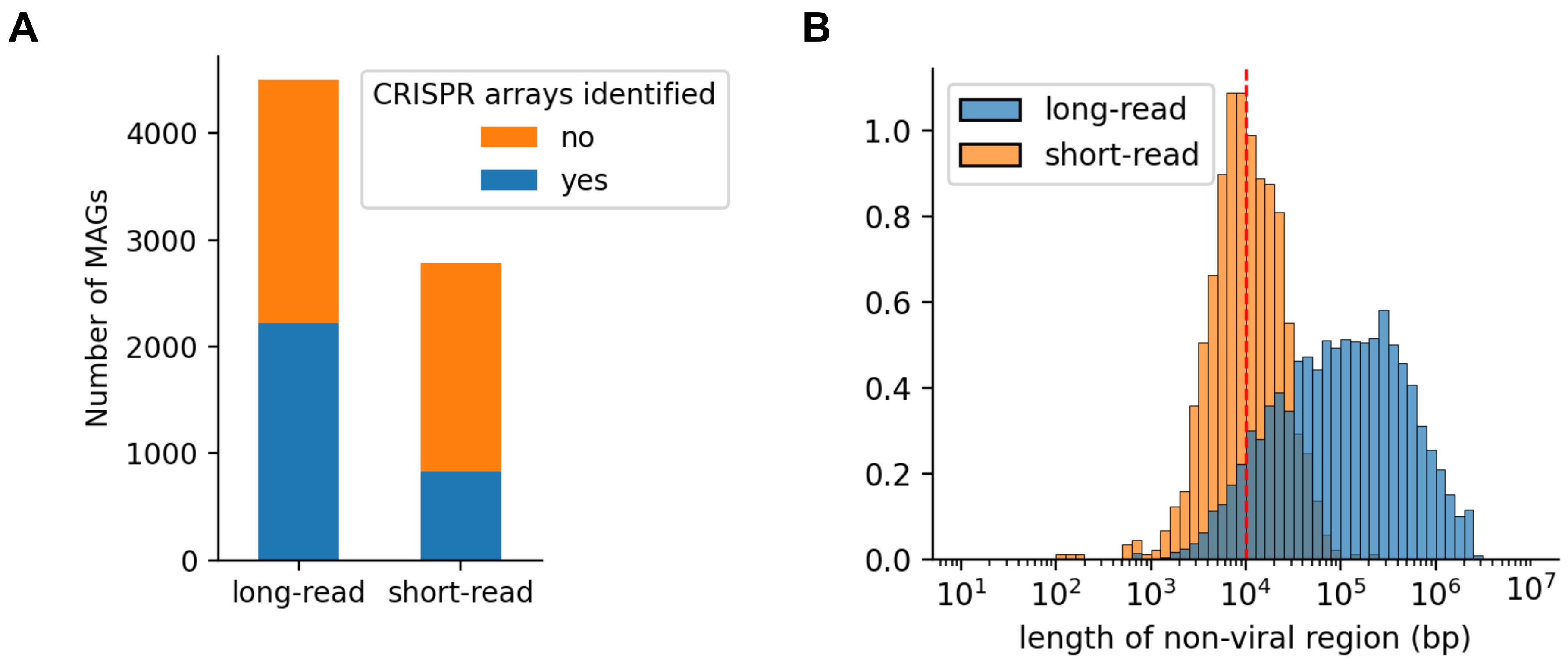


<u>**Supplementary Figure 13:**</u> **Long-read metagenomic assemblies improve phage-host linkage analysis.** (A) The number of long-read–augmented and short-read–only MAGs with CRISPR arrays identified (blue) hence enabling host association via spacer matching. (B) Length distributions of flanking non-viral regions surrounding putative prophages based on long-read–augmented (blue) and short-read–only (orange) metagenomic assemblies. Prophages with at least 10kbp of flanking non-viral regions (red dotted line) were used for host association, to avoid linking contigs that are almost entirely viral (which are more likely to be incorrectly binned).

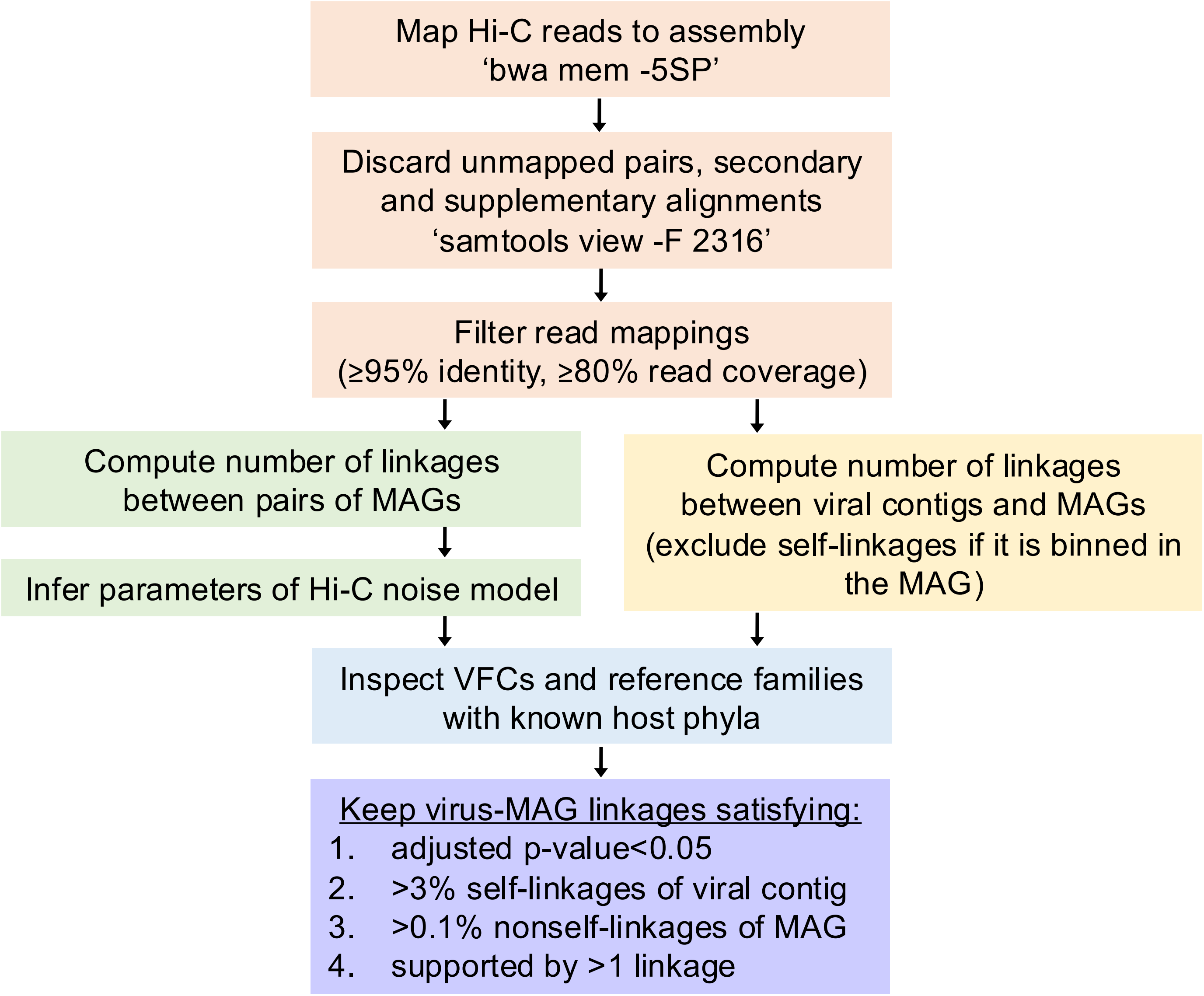


**Supplementary Figure 14: Hi-C host association pipeline.** A mathematical model for noise in Hi-C data was constructed based on the assumption that most Hi-C linkages between MAGs are from non-specific ligation in extracellular DNA. Parameters for each MAG's contribution to this noise were inferred on a per-sample basis. This was used to estimate a p-value for the linkages between a viral contig and a MAG. Further filtering was applied to reduce potential false positives (**Methods**).

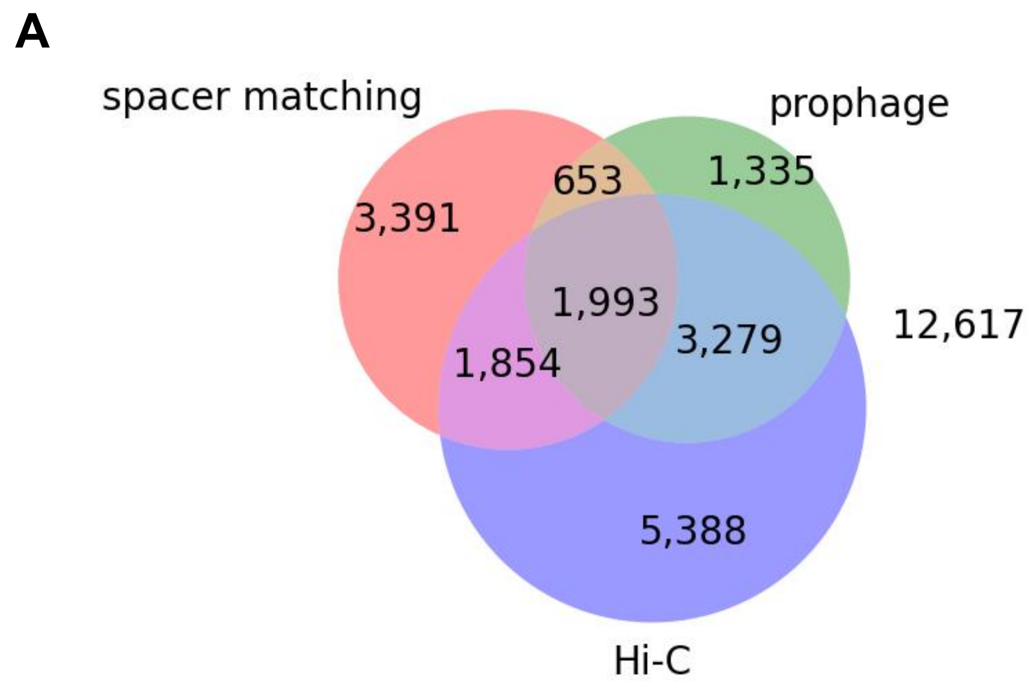


**<u>Supplementary Figure 15:</u> Comparison of information provided by host association methods.** (A) Number of vOTUs assigned a host by each method. Hi-C assigned hosts to 41%, spacer matching to 26%, and prophage analysis to 24% of vOTUs. (B) Comparison of host assignments at the VFC level, showing greater shared assignments across methods.

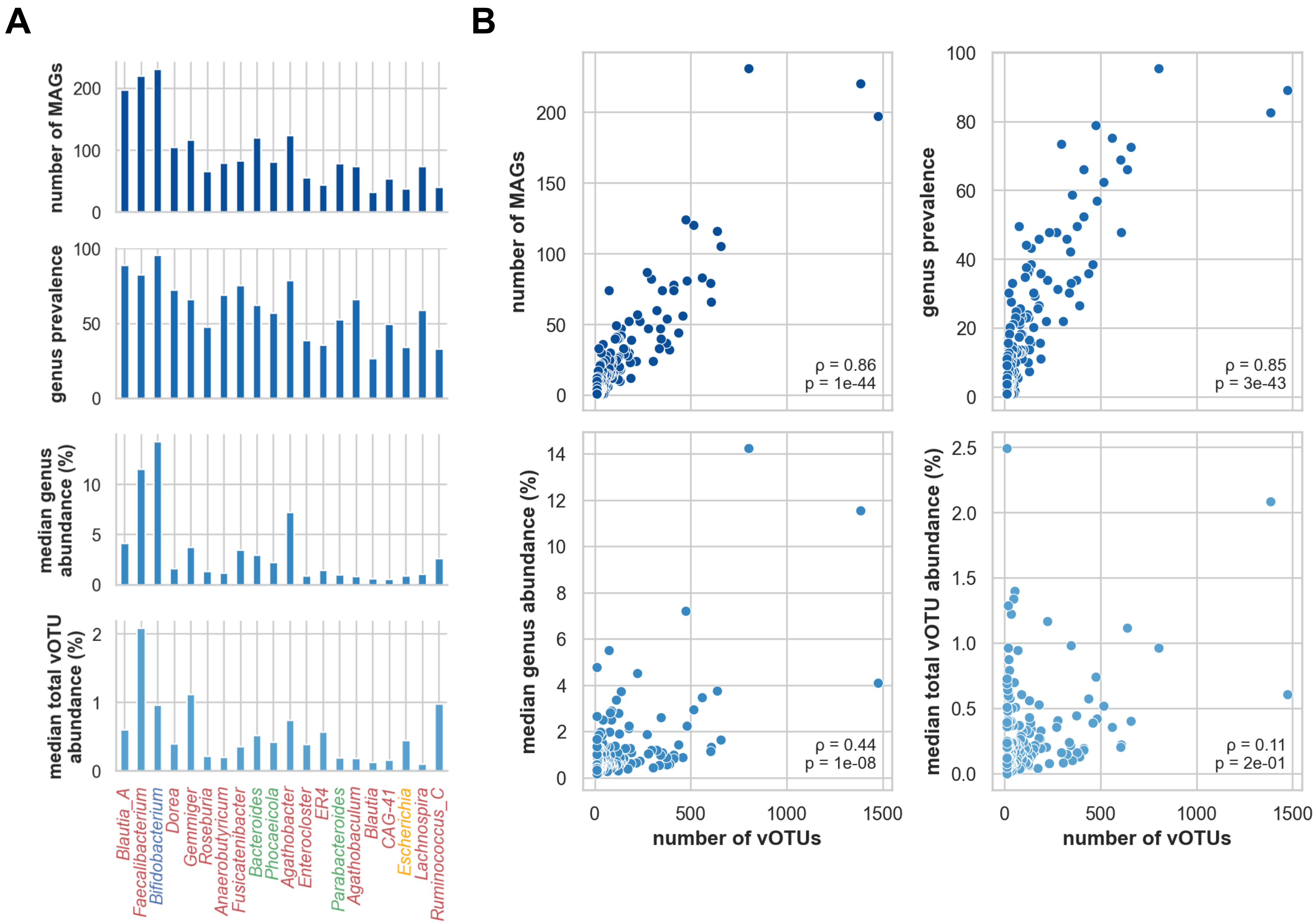


**<u>Supplementary Figure 16:</u> vOTU diversity correlates with the number of MAGs, prevalence, and median abundance of host genera.** (A) The number of MAGs assembled, prevalence, median abundance, and median vOTU abundance of host genera, ordered by the number of vOTUs assigned (**Fig. 2B**). The top 20 genera are shown. (B) Correlation plots between each feature and the number of vOTUs, for genera with ≥10 vOTUs assigned (n=151). Spearman correlation coefficients and p-values are shown in the lower right of each plot.

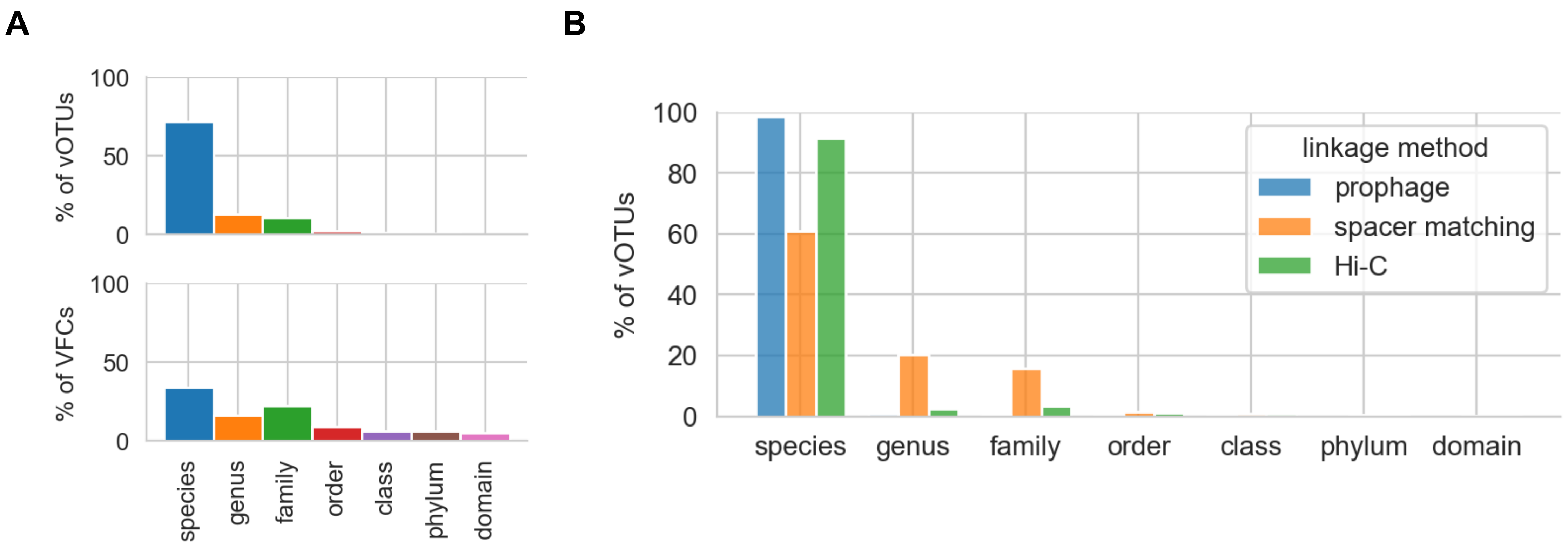


**<u>Supplementary Figure 17:</u> Host range specificity distributions of vOTUs and VFCs.** (A) Distribution of host range specificity of vOTUs (upper) and VFCs (lower). 15% of vOTUs, and 50% of VFCs, infect >1 genus. (B) Stratifying by host linkage method, spacer matching identifies the broadest host ranges of vOTUs, with 19% linking to >1 genus.

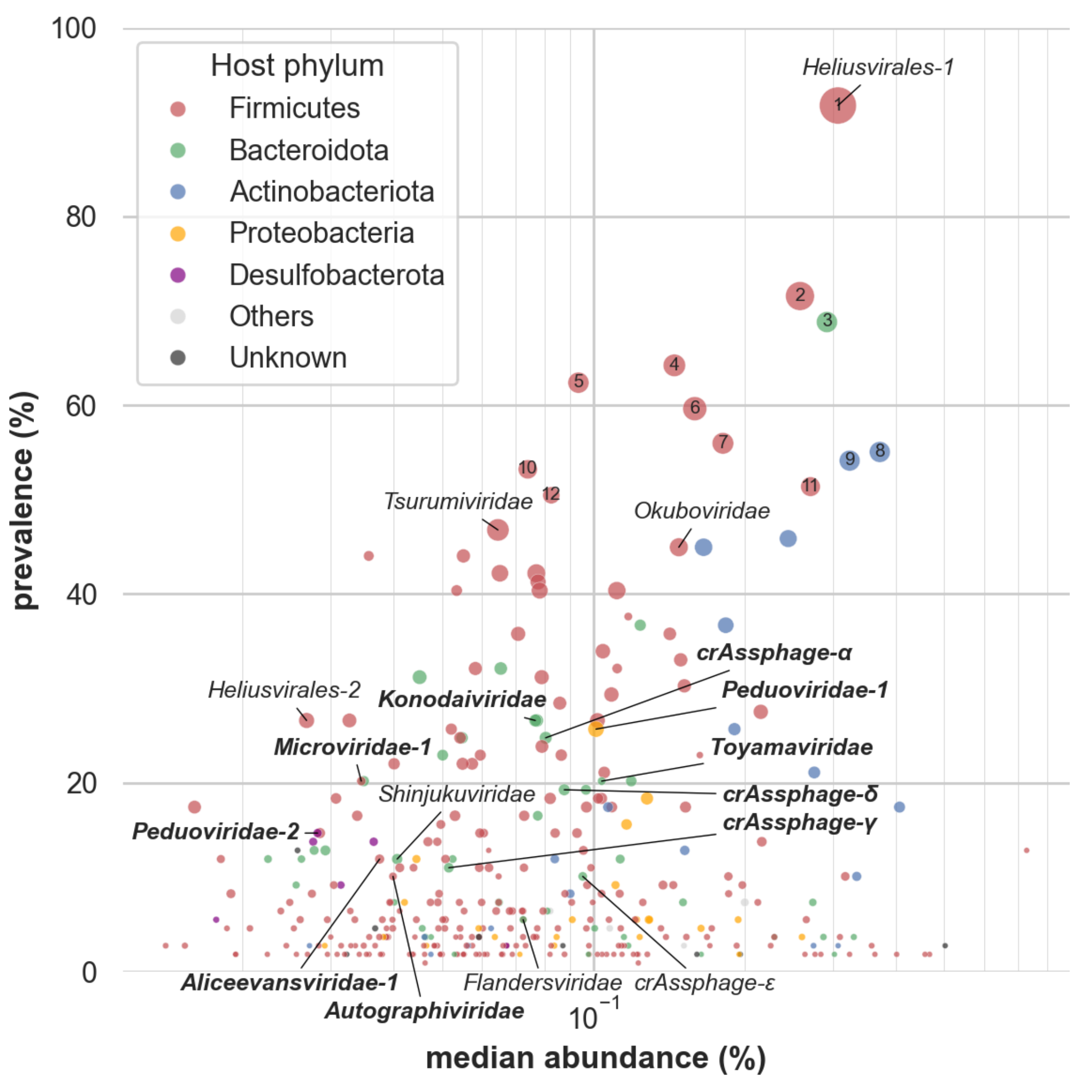


**<u>Supplementary Figure 18:</u> Primary host phyla of the VFCs.** Prevalence versus median abundance of the VFCs (**Fig. 1D**) color-coded by host phylum. ICTV-ratified families are highlighted in bold.

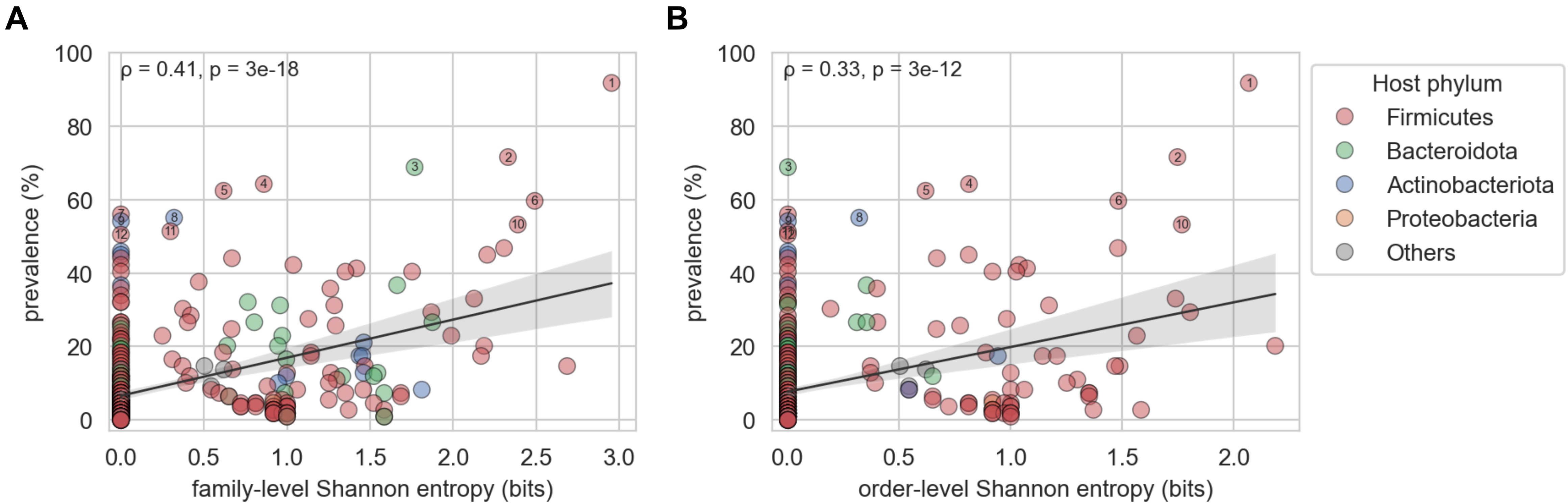


**<u>Supplementary Figure 19:</u> VFC host range and prevalence are correlated.** Scatterplot depicting entropy of host taxa at the family (A) and order (B) level for VFCs versus prevalence in the SPMP cohort. VFCs are color-coded by host phylum, and those with >50% abundance are labelled (1-12). Spearman correlation coefficients and p-values are shown in the upper left of each plot.

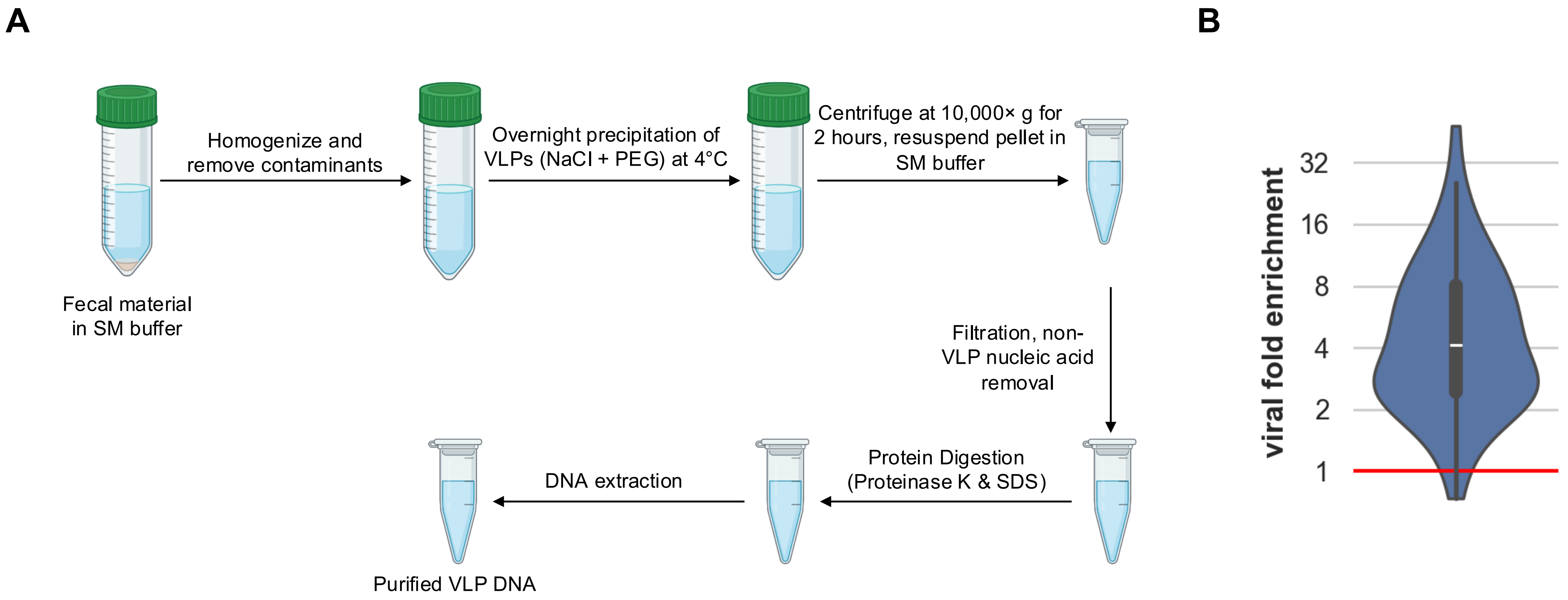


**<u>Supplementary Figure 20:</u> Virus-like particle enrichment from stool samples.** (A) Schematic of the virus-like particle (VLP) enrichment protocol (**Methods**; created with BioRender.com). (B) Degree of viral enrichment as quantified by the ratio of 16S SSU rRNA mapping rates to the bulk versus VLP samples. A ratio of 1 (red line) indicates no viral enrichment. The VLP samples (n=64) exhibit high viral enrichment with respect to the bulk (median fold enrichment of 4.1).

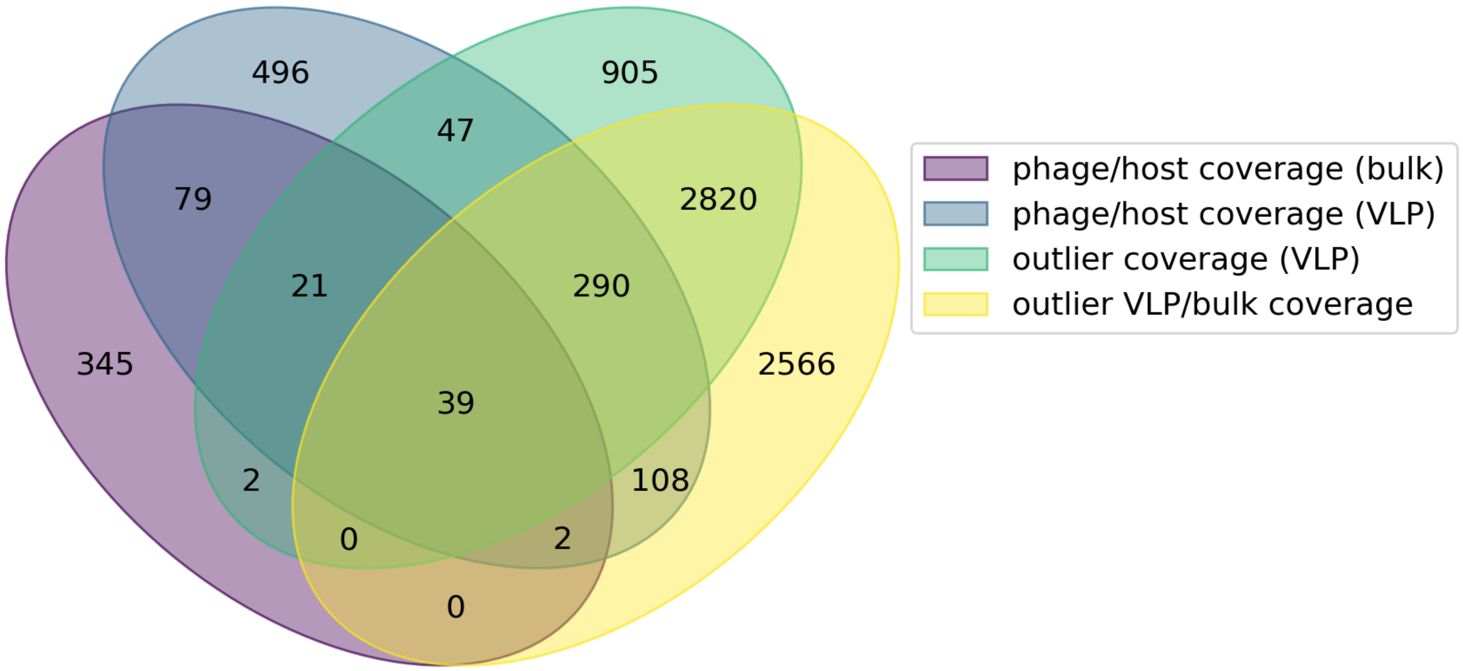


**<u>Supplementary Figure 21:</u> Identification of active replication signatures for thousands of vOTUs.** Venn diagram of vOTU-sample pairs satisfying each of the following metrics: 1) vOTU/host MAG mean coverage in bulk metagenome >1.75 [phage/host coverage (bulk)]; 2) vOTU/host MAG mean coverage in VLP metagenome >1.75 [phage/host coverage (VLP)]; 3) vOTU mean coverage in VLP metagenome exceeds outlier threshold based on non-viral contigs [outlier coverage (VLP)]; 4) VLP/bulk RPKM ratio exceeds outlier threshold based on non-viral contigs ([outlier VLP/bulk coverage]; **Methods**; **Supplementary Data 7-8**). >6,000 vOTUs were detected to be replicating by at least one measure. All pairs of measures have non-random overlaps (two-sided Fisher's exact test with FDR correction).

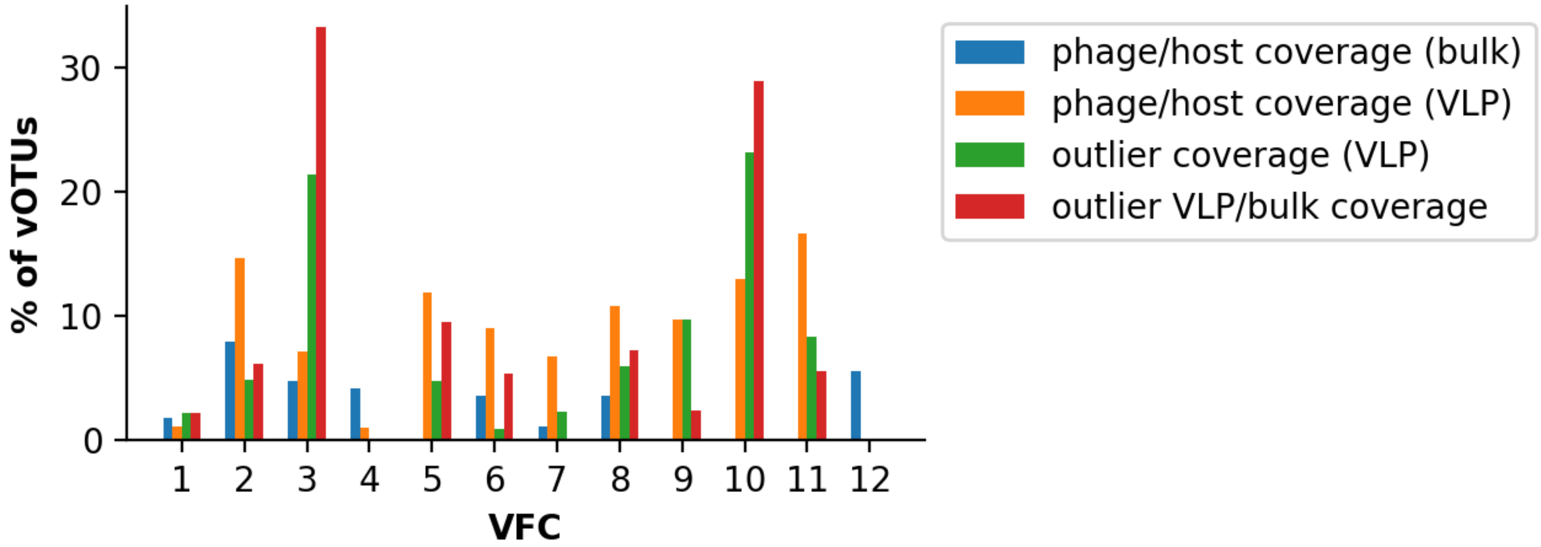


**Supplementary Figure 22: Active replication signatures of the top VFCs.** Percentage of vOTUs in the most prevalent VFCs satisfying each of the active replication detection approaches (**Supplementary Fig. 21**) in at least one sample.

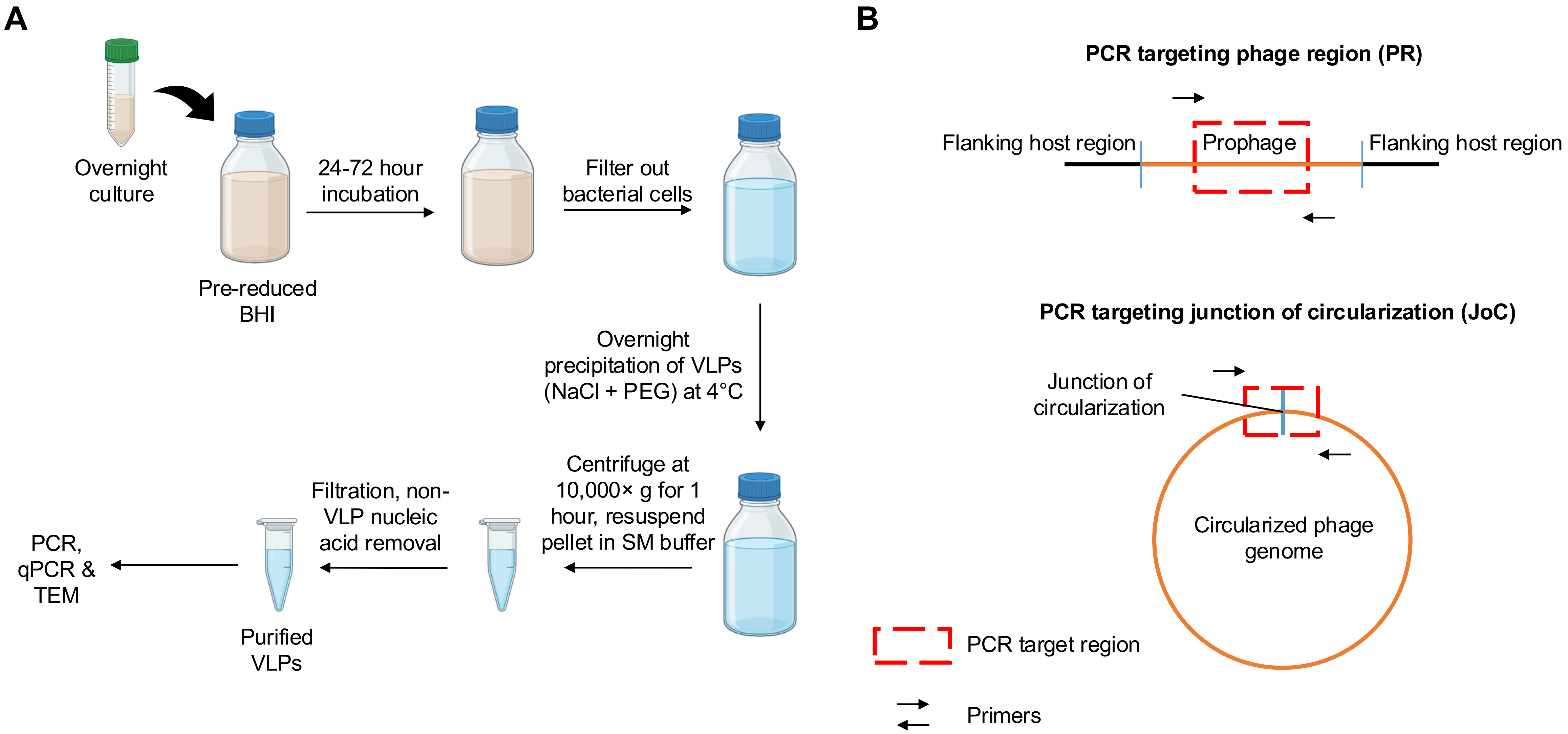


**<u>Supplementary Figure 23:</u> Experimental workflow for prophage induction and purification of induced VLPs.** (A) Schematic of the protocol to induce prophages from bacterial cultures and purify induced VLPs (**Methods**; created with BioRender.com). (B) Schematic of primers designed to target the internal phage region (PR) and junction of circularization (JoC) of circularized phage.

**<u>Supplementary Table 1:</u> Bacterial strains and prophage locations**

| Strain name | ATCC | Family | VFC | Contig ID | Prophage coordinates (geNomad prediction) | Prophage coordinates (replicating region) | Comments |
|---|---|---|---|---|---|---|---|
| *Ruminococcus torques* Holdeman and Moore | 22756 | *Lachnospiraceae* | 1 | a1883c1a3df8477f_1 | 1693947-1742354 | 1692600-1748695 | |
| *Ruminococcus gnavus* Moore et al. | 29149 | *Lachnospiraceae* | 2 | NZ_CP027002.1 | 1389840-1434063 | 1389680-1435651 | |
| *Anaerostipes hadrus* (ATCC 29173) | 29173 | *Lachnospiraceae* | 4 | 72d052e7488c4515_3 | 82-43705 | 1-39447 | Prophage on 72d052e7488c4515_4, 32139-41020 is part of the same phage |
| *Agathobacter rectalis* (Hauduroy et al.) Prevot | 33656 | *Lachnospiraceae* | 4 | NC_012781.1 | 1928248-1993269 | 1928876-1993526 | |

**Supplementary Table 2:** **Full-length 16S primers**

| Primer | Primer sequence |
| --- | --- |
| forward primer (S-D-Bact-0008-c-S-20) | AGRGTTYGATYMTGGCTCAG |
| reverse primer (1492R) | CGGYTACCTTGTTACGACTT |

**Supplementary Table 3:** **Primers used in PCR-based validation of phage induction**

| Bacterial strain | Target region | Primer | Primer sequence | Annealing temperature used (°C) |
|---|---|---|---|---|
| *Agathobacter rectalis* ATCC 33656 | Phage region (PR) | AR_p1-F | CATCTTCCTGCACCTCCTCTG | 56.3 |
| | | AR_p1-R | CAGACATTTCGGAGTATTGCTGCG | |
| | Junction of circularization (JoC) | AR_pL | TACGCCAGGGAATTATCCTTG | 51.8 |
| | | AR_pR | TCGTGGTTGCGGTATTTC | |
| *Anaerostipes hadrus* ATCC 29173 | Phage region (PR) | AH_p1-F | GGATTACCATGCTGATGGAAGAGG | 56.3 |
| | | AH_p1-R | GCTGGTATCACAGTAACTGCACC | |
| | Junction of circularization (JoC) | AH_pL3 | GATGCAAAAGTAAATGAATATGCTC | 53.7 |
| | | AH_pR3 | GAGCGACAGAGCTTGTAATTG | |
| *Ruminococcus gnavus* ATCC 29149 | Phage region (PR) | RG_p2-F | CGGGCCTTCGATGTTCTCATAG | 54.7 |
| | | RG_p2-R | CCTTCGTTCCGGCAAAATACAC | |
| | Junction of circularization (JoC) | RG_pL | CATACAACGGAGTGAAGTGG | 50.8 |
| | | RG_pR | GTGATACATATAAAAGGGATTATTCCG | |
| *Ruminococcus torques* ATCC 27756 | Phage region (PR) | RT_p1-F | CCAAGCATGTCCAGAACTCCAAG | 57.1 |
| | | RT_p1-R | GCACAGGTGCTTTAATGGGCG | |
| | Junction of circularization (JoC) | RT_pL | TGGAAAGAATTAGTAAAAAGGATAGG | 50.1 |
| | | RT_pR | CATATTCGTCTTCTCCTTTTGCC | |

**Supplementary Table 4: Primers used in qPCR**

| Bacterial strain | G-Block sequence | Primer | Primer sequence | Annealing temperature used (°C) |
|---|---|---|---|---|
| *Agathobacter rectalis* ATCC 33656 | GTCCAAATCCTGCACTTCACTTTCTGTCAGCTCTC<br>CAATCCACTCTCCAATTCTTTCCTCCGAAACCGTG<br>GAAATCTGCTCGCACAAGAGTGTTGATGGTCTAAG<br>TGCTGACTCAATATATACATGAGTCGGAAGGTCAG<br>TCTTTGGTTTGGTCG | 2q01_AR-Fwd | GTCCAAATCCTGCACTTCAC | 52.9 |
| | | 2q01_AR-Rev | CGACCAAACCAAAGACTGAC | |
| *Anaerostipes hadrus* ATCC 29173 | TTGAAGATGTGAAAGGCGTGGAAACAGATGTTTTC<br>AAGATCAAGAAAAAGATGTTCAAAAAGAAATATGG<br>AGATCTGTACGATTTACGAATAACGAGGTGATCAC<br>ATGAAGCAAAAGAGCAGCTTCCTGATCTACCATGA<br>ATATCGGGAACCACTAAAATTACTGACAGATGAGC<br>AGAGAGGTCGGTTATTGATGGCAT | 2q02_AH2-Fwd | TTGAAGATGTGAAAGGCGTG | 52.1 |
| | | 2q02_AH2-Rev | ATGCCATCAATAACCGACCTC | |
| *Ruminococcus gnavus* ATCC 29149 | GATCAGCACATGGTTCATCGGGTCATCGACCTTCT<br>CGAAATAGGTCTGAATGAAATTGTTGCCGTGCCAG<br>TTAAACATTCTTCTGACTGCGAGCCAGATGTCTTTT<br>GCATCTCCACTGGAAGGAAATGCACCTGTGTAGTT<br>GCCCCACAGCTTCCATCCATTCATGTTGATTGCTG<br>TTGCTACTCCATAGGT | q02_RG-Fwd | GATCAGCACATGGTTCATCG | 52.8 |
| | | q02_RG-Rev | ACCTATGGAGTAGCAACAGC | |
| *Ruminococcus torques* ATCC 27756 | ACACTTTGAGCAGATTCCCATCAGGAATCTTGTAT<br>CCAACCAGGAATATCAGAGAAATCTTTCACAGAAT<br>CATGTCCAGCGTGCCGCTGCGAACTTCGATCTGT<br>ACCAGATAAATCCCGTAAAAGTCAGCCGCCGTAAC<br>GGCATCAACTATGTCTTTAACGGGCAGCACACCAT<br>TGAAATCGT | q01_RT-Fwd | ACACTTTGAGCAGATTCCC | 51.5 |
| | | q01_RT-Rev | ACGATTTCAATGGTGTGCTG | |

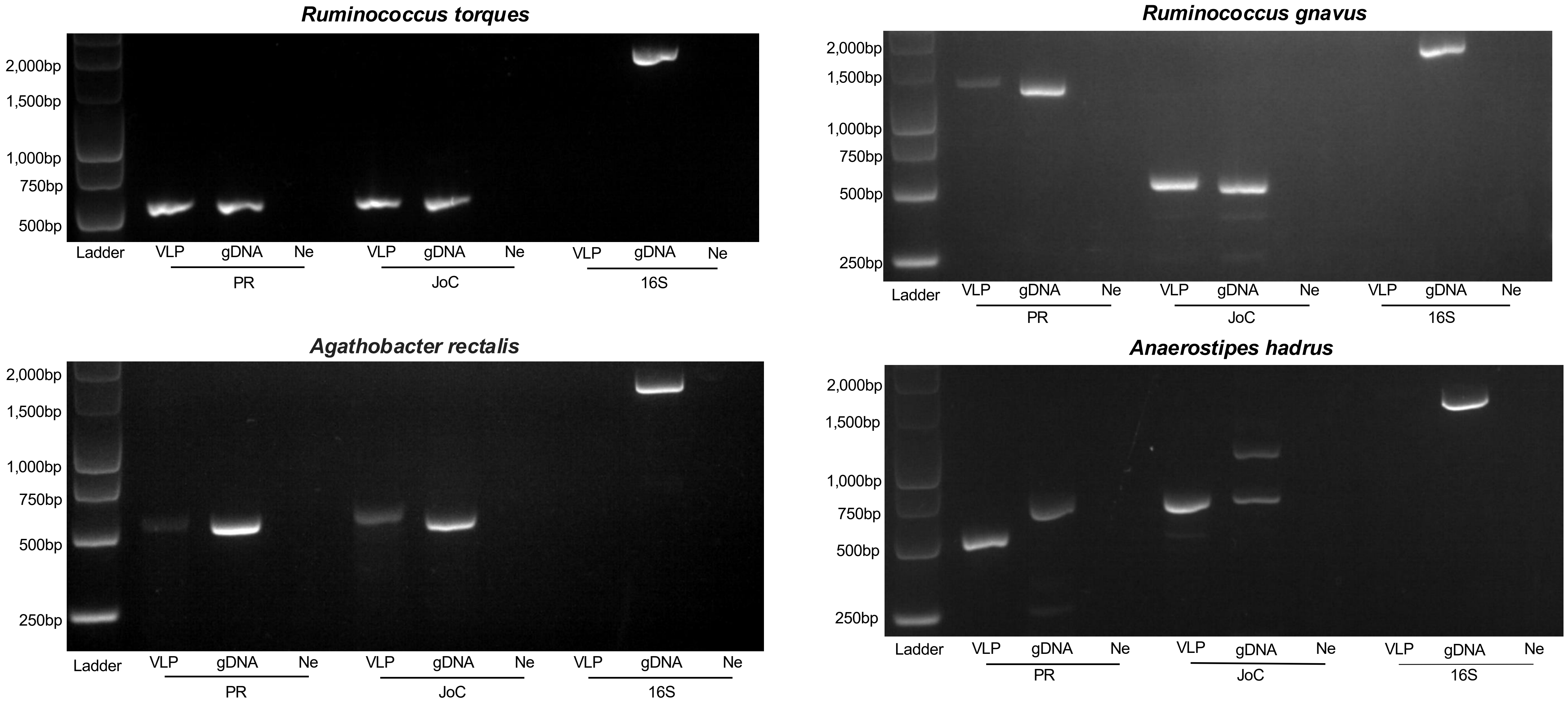


**<u>Supplementary Figure 24:</u> PCR evidence for prophage induction and virion formation in four bacterial hosts.** Two sets of PCR primers were designed for VFC 1-4 prophages in *Ruminococcus torques*, *Ruminococcus gnavus*, *Anaerostipes hadrus*, and *Agathobacter rectalis* (**Supplementary Table 3**): one in the internal phage region (PR) and one spanning the prophage ends testing for circularized DNA (junction of circularization, JoC). PCR bands were observed for the JoC primer in genomic DNA (gDNA) for all four isolates, indicating that all phages induced and were replicating. All phages also have PCR bands for both primers in VLP-purified DNA, indicating formation of complete phage particles. 16S bands in gDNA but not VLP DNA for all isolates confirmed the lack of bacterial contamination in the VLP fraction.

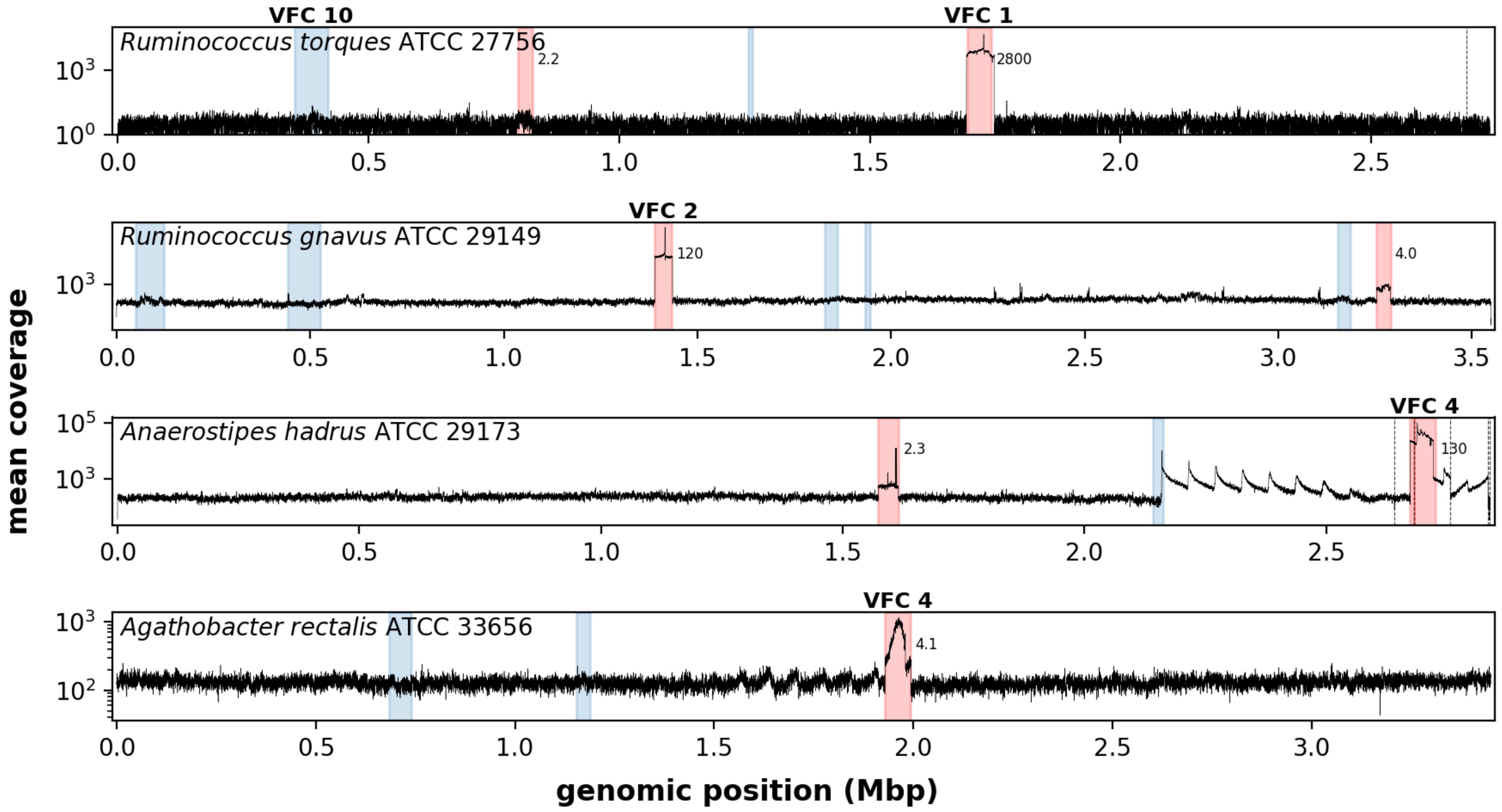


**<u>Supplementary Figure 25:</u> Prophages from the top VFCs spontaneously induce.** Coverage depth of VLP sequencing data from the no mitC added condition for the same strains as in **Fig. 3C**. Vertical dotted lines indicate contig boundaries. Blue and red shaded regions denote geNomad-predicted prophages. Regions shaded in red have mean coverage >2× of the host, with mean coverage ratios indicated.

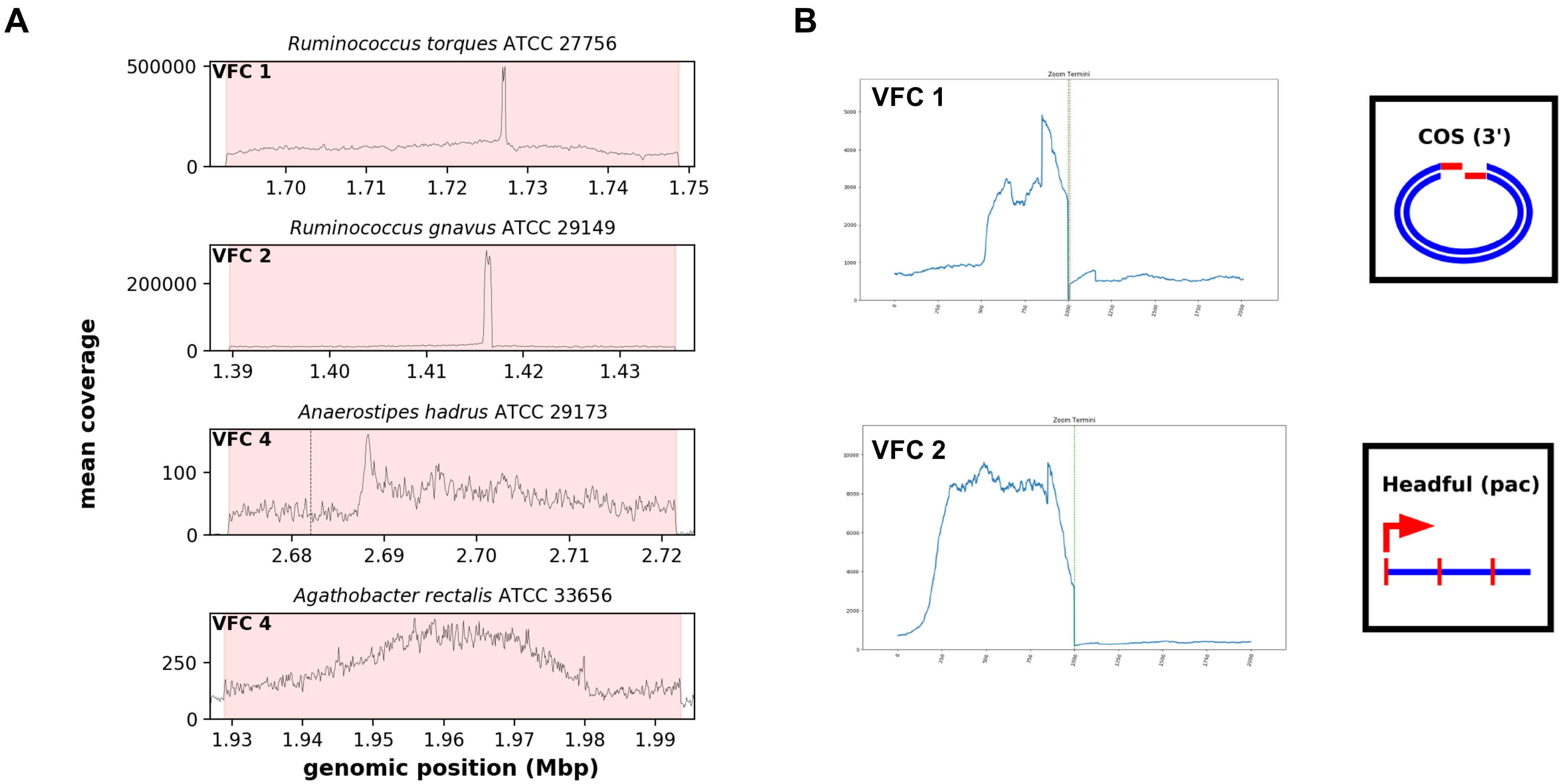


**<u>Supplementary Figure 26:</u> Replication strategies of induced prophages.** (A) Mean coverage in the four VFC 1-4 prophage regions with the observed replicating regions highlighted and plotted on a linear scale (zoomed-in version of **Fig. 3D**). (B) PhageTerm analysis indicates that the VFC 1 and 2 phages utilize a cos (3') and headful packaging mechanism, respectively, while results for the VFC 4 phages were inconclusive.

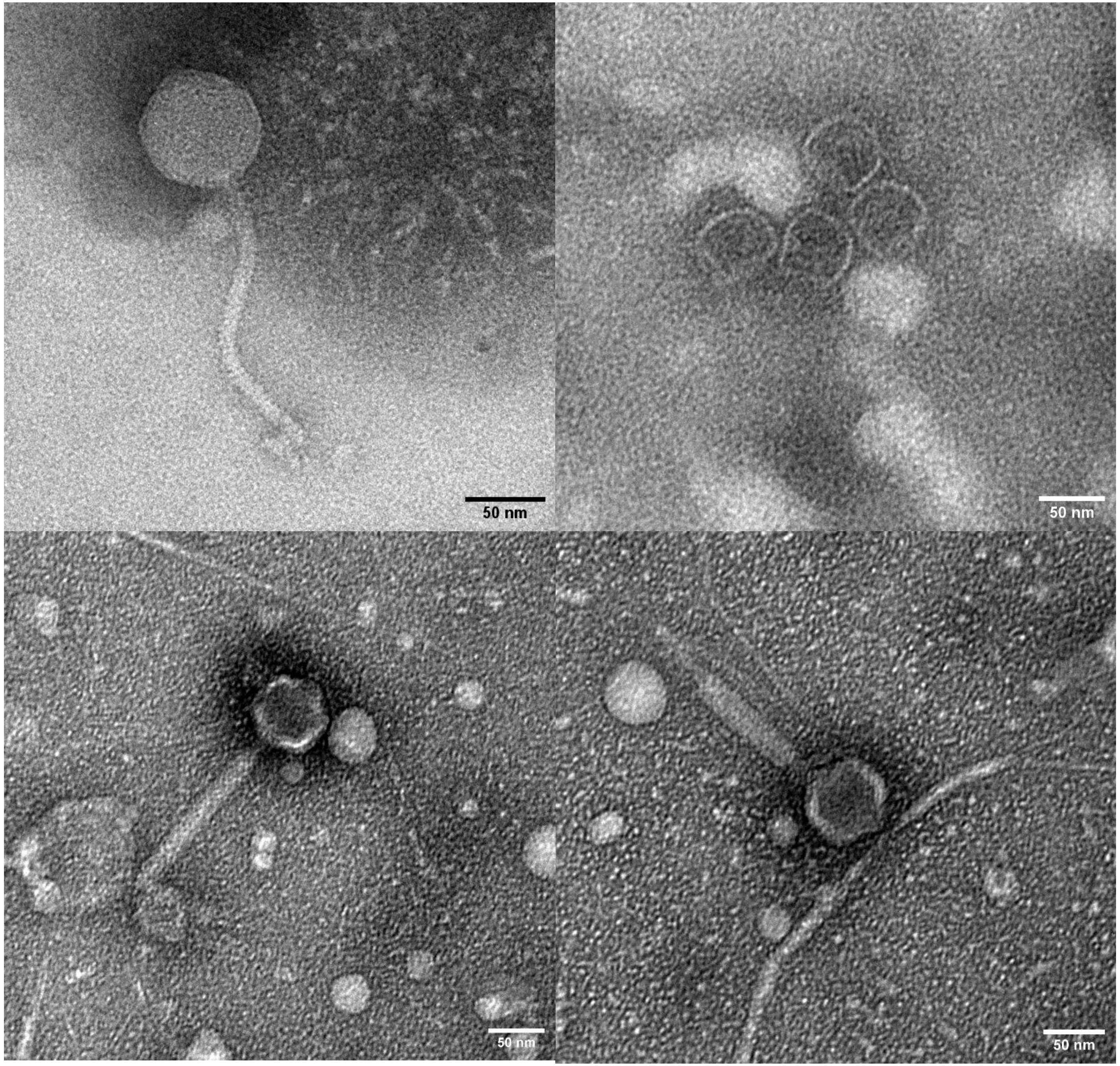


**<u>Supplementary Figure 27:</u> TEM images of induced phages.** These images present a larger field of view for the images shown in **Fig. 3E**.

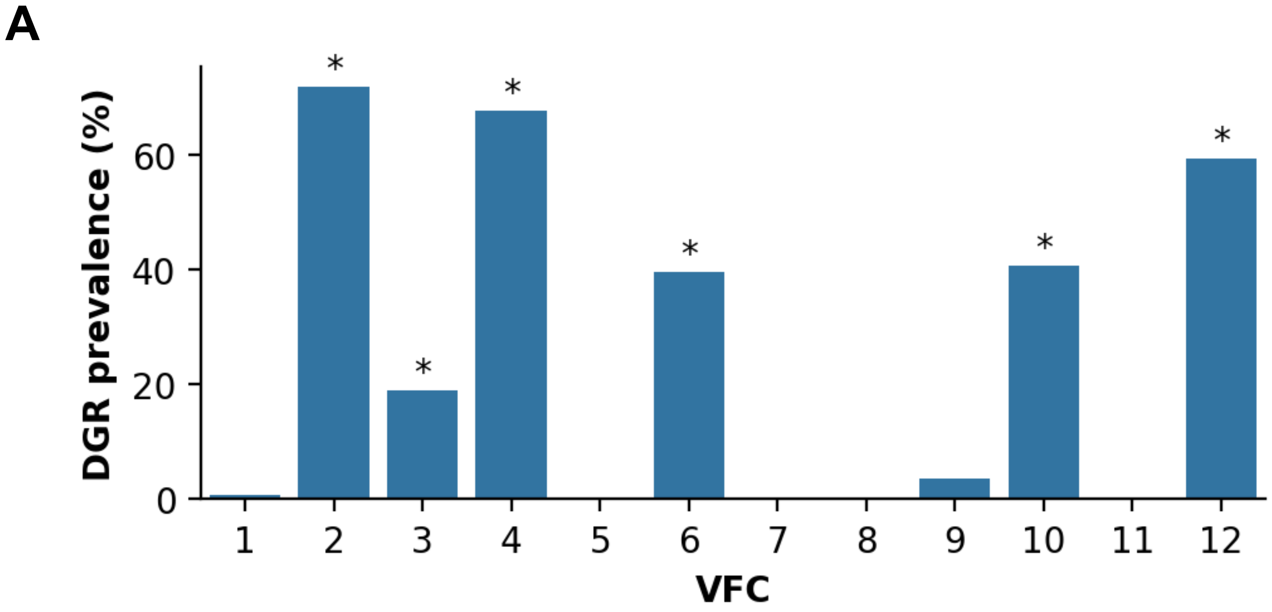


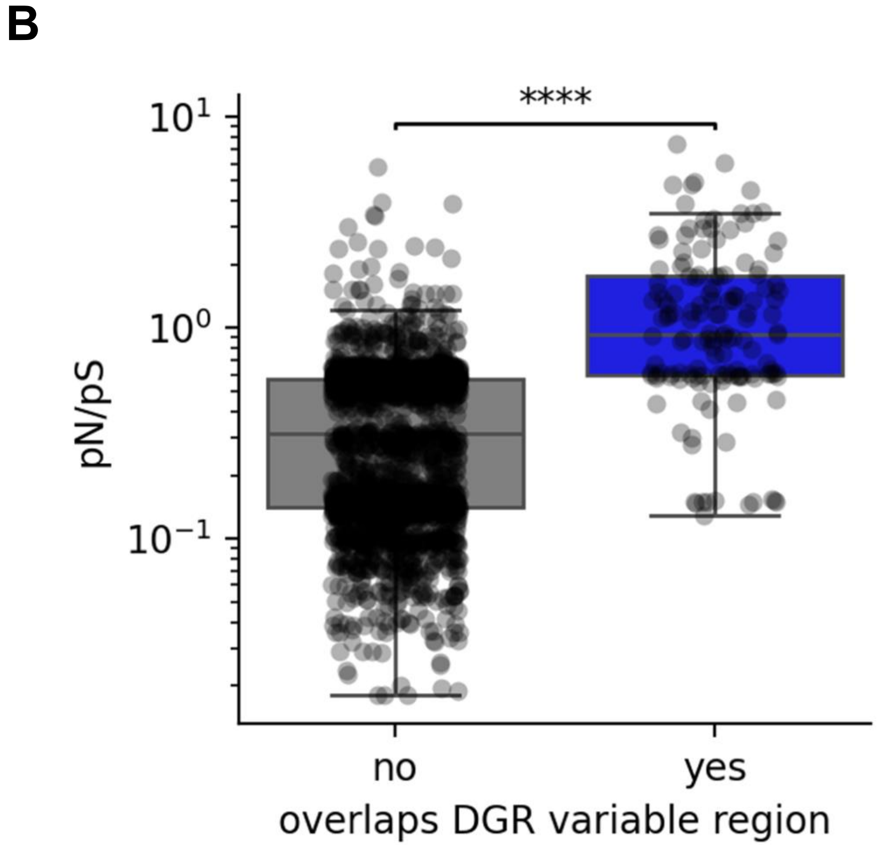


C

| Gene annotation | No. of genes |
|---|---|
| Putative phage holin Dp-1 | 14 |
| Bacteriophage lambda head decoration protein D | 10 |
| Phage-related tail fibre protein | 10 |
| CotH kinase protein | 9 |
| BppU N-terminal domain | 5 |

**Supplementary Figure 28: Several highly prevalent VFCs utilize diversity generating retroelements (DGRs) for host range diversification and immune evasion.** (A) Prevalence of DGRs in the top VFCs. VFCs whose DGR prevalence are outliers (>median+2×IQR) relative to all VFCs are indicated with a star. (B) Boxplots depicting the pN/pS ratio for genes stratified by the overlap with DGR variable regions. Genes containing DGR variable regions show significantly higher pN/pS ratios (median 0.9 vs 0.3; two-sided Mann-Whitney U test p-value<$10^{-6}$). (C) The most frequent annotations of genes in the top VFCs overlapping with DGR variable regions.

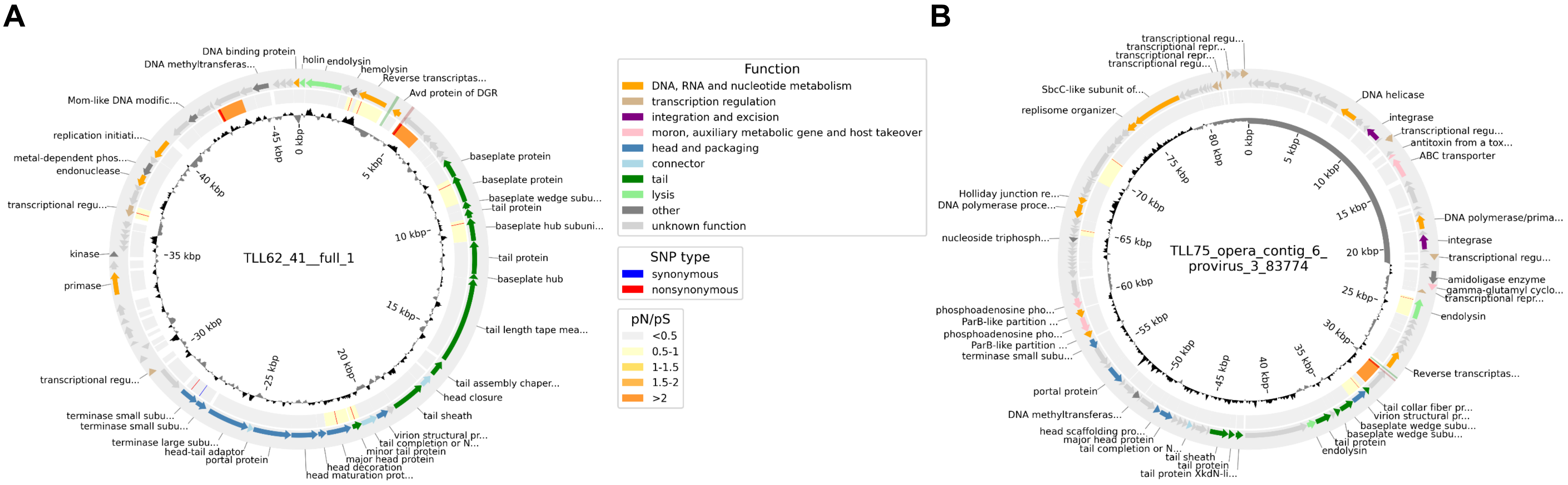


**Supplementary Figure 29: Active DGRs are associated with host range switching.** Circos plots of vOTUs carrying active DGRs. The outer track shows gene annotations color-coded by PHROG category; the middle track shows synonymous (blue) and nonsynonymous (red) SNPs, and a heat map of pN/pS ratios of genes (**Methods**); and the inner track shows coverage skew. DGR template (green) and variable (red) regions are highlighted spanning the outer and middle tracks. (A) A VFC 2 vOTU was linked to 2 *Gemmiger* species by Hi-C. A few nonsynonymous SNPs were detected, most prominently in two DGR variable regions (one undetected by DGRscan). (B) A VFC 4 vOTU was linked to *Mediterraneibacter faecis* and *Fusicatenibacter saccharivorans* by Hi-C. A few nonsynonymous SNPs were detected, mostly in one DGR variable region.

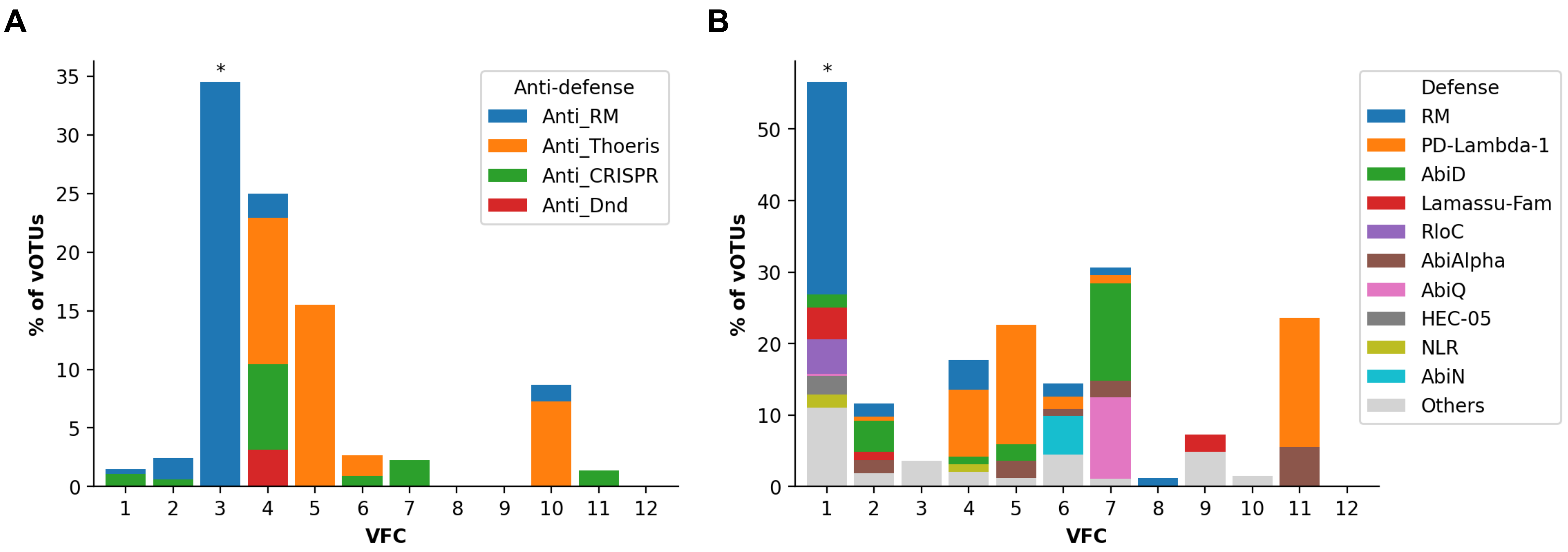


**<u>Supplementary Figure 30:</u> The top VFCs carry diverse systems for phage defense and anti-defense.** (A) Distribution of anti-antiphage defense systems in VFCs 1-12. (B) Distribution of antiphage defense systems in VFCs 1-12. VFCs whose defense and anti-defense prevalence are outliers (>median+2×IQR) relative to all VFCs are indicated with a star.

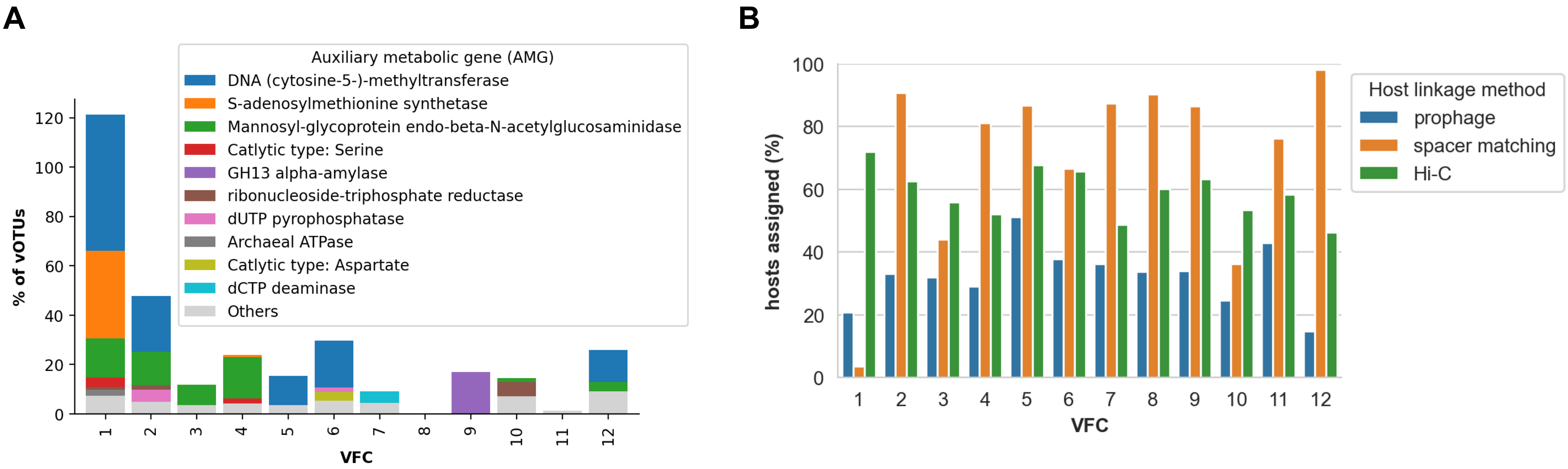


**Supplementary Figure 31: Methyltransferases may mediate anti-CRISPR defense in VFC 1.** (A) Distribution of auxiliary metabolic gene (AMG) functions across the top VFCs. VFC 1 phages frequently carry DNA methyltransferases and SAM synthetases, which are involved in DNA modification, protecting the phage from host immune defences. (B) Distribution of host assignments by host linkage method in the top VFCs. vOTUs in VFC 1 have very low spacer matching occurrence. As known anti-CRISPR systems were not detected, we hypothesize that VFC 1 either carries a novel anti-CRISPR system, or there is a DNA-modification-based anti-CRISPR function.

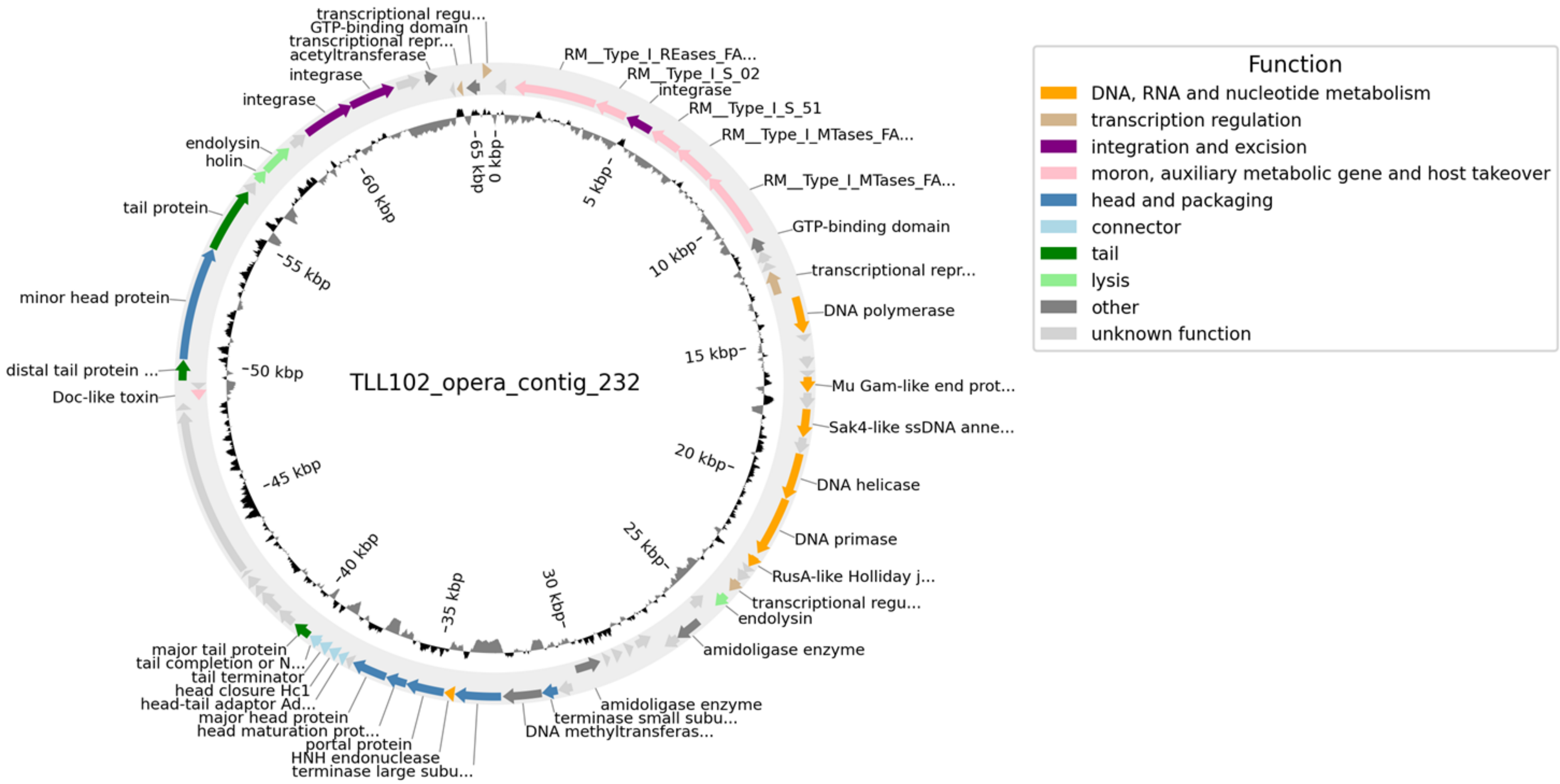


**<u>Supplementary Figure 32:</u> A VFC 1 vOTU with broad host range.** Circos plot of a VFC 1 vOTU carrying a restriction-modification system and linked to *RUG472 sp900545265* and *UMGS1603* by Hi-C. The outer track shows gene annotations color-coded by PHROG category; and the inner track shows coverage skew.

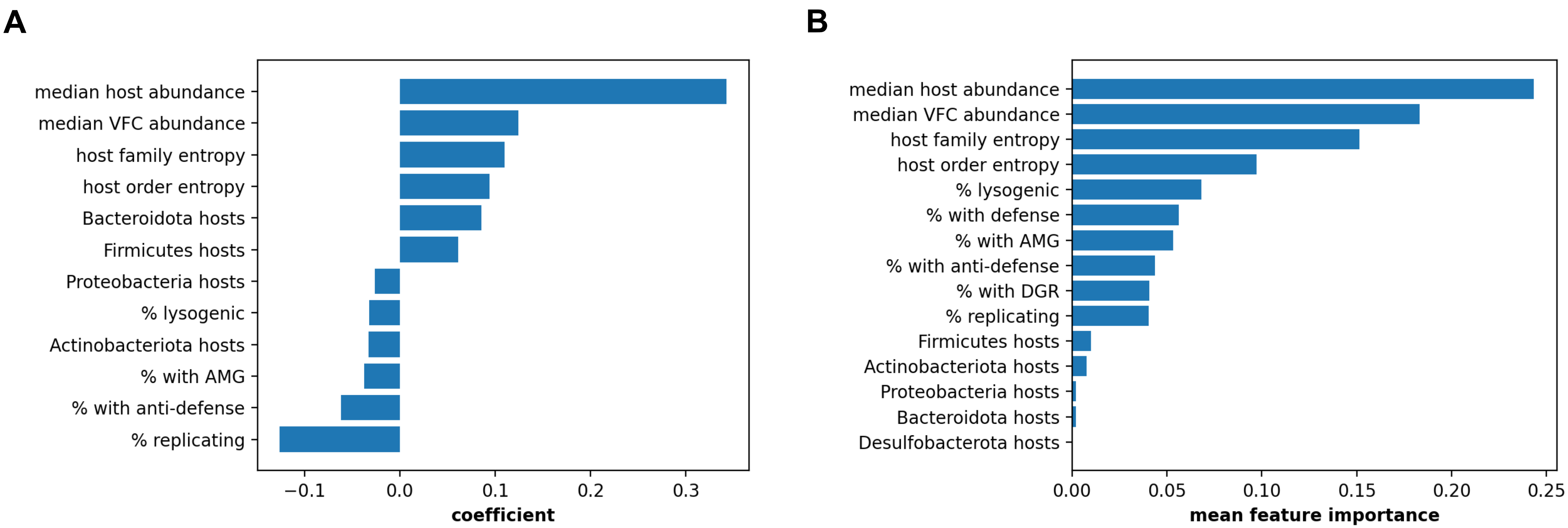


**Supplementary Figure 33: Host abundance and host range breadth predict VFC prevalence.** (A) A generalized linear model was constructed to predict VFC prevalence in the SPMP cohort (**Methods**). VFCs with size ≥10 were used (n=120). The fit explained 48% of the variance. (B) A random forest regression model was constructed on the same set of VFCs (**Methods**). The model explained 21% of the variance.
